# Character displacement within the breeding area questions reinforcement in *Ficedula* flycatchers

**DOI:** 10.1101/515916

**Authors:** Vladimir G. Grinkov, Igor V. Palko, Helmut Sternberg

## Abstract

At present, studies of reinforcement should be focused on demonstrating how often this process occurs in nature and how important it is for speciation. Here we study the character displacement within the breeding area in the Pied Flycatcher to check the validity of the reinforcement in *Ficedula* flycatchers. We used point-referenced spatial data and a random forest to find the most important explanatory factors of the character displacement, and to reconstruct the phenotypic structure of the populations. The environmental temperature, and not the distance to sympatry, were proven to better describe the geographic pattern of the mean breeding plumage colour of the Pied Flycatcher populations. We conclude that ecologically distinct adaptations drive the morphological differentiation of the Old World flycatchers, and not reinforcement.

## Introduction

Reinforcement (Dobzhansky 1937) is the process of enhancing reproductive isolation directly controlled by natural selection to cancel maladaptive hybridization between nascent species living together in the same territory (in sympatry). Often reinforcement is considered as one of the ways to complete the speciation process (Lewontin 1974; Servedio 2004). Scientific acceptance of the theory at the time of its appearance was very high, but in the eighties, it fell drastically (Noor 1999). To date, however, doubts about the existence of reinforcement have been resolved both by theoretical studies and by the identification of such cases in nature (Noor 1999; Servedio and Noor 2003; Butlin and Smadja 2018). Now researchers would change the focus of reinforcement studies to show how often this process occurs in nature and how important it is for speciation (Servedio and Noor 2003). Therefore, we are addressing to answer these questions by the case study of the character displacement (Brown and Wilson 1956) within the breeding area in the Pied Flycatcher (*Ficedula hypoleuca*).

After the publication of Sætre et al. (Saetre et al. 1997), relationships between the Pied Flycatcher and the Collared Flycatcher (*F. albicollis*) had come to be considered one of the examples of reinforcement, although similar ideas were expressed several times before (Kral et al. 1988; Alatalo et al. 1990a; Lundberg and Alatalo 1992; Alatalo et al. 1994). The Pied Flycatcher breeding range is many times larger than the one of the Collared Flycatcher (Fig. 1). Most of the Pied Flycatcher breeding area does not overlap with the one of the Collared Flycatcher (allopatric populations), while most of the Collared Flycatcher breeding range coincides with the breeding area of the Pied Flycatcher (Fig. 1). Where the ranges of these two species of flycatchers overlap (areas of sympatry, for example, in Central and Eastern Europe), males with a brown colouration of breeding plumage (brown morph) predominate in populations of the Pied Flycatcher. This colouration is very similar to that of females of this species. In the allopatric area (for example, in Fennoscandia), contrastingly coloured males with a black and white colour of the breeding plumage (black morph) predominate in populations of the Pied Flycatcher (Fig. 1 A and B). Such a biographical pattern of the morph ratio in the Pied Flycatcher is fundamental to the assumption that the character displacement within the breeding area of the species is the result of reinforcement (Saetre et al. 1997). However, the coupling of a high proportion of brown males with areas of sympatry in the Pied Flycatcher is revealed only for European populations, without analysing the phenotypic structure of those populations that breed in the most eastern European and the Asian part of the species breeding range (for example, in the European part of Russia to the Ural Mountains and further to Western Siberia, Fig. 1C).

**Fig. 1.**
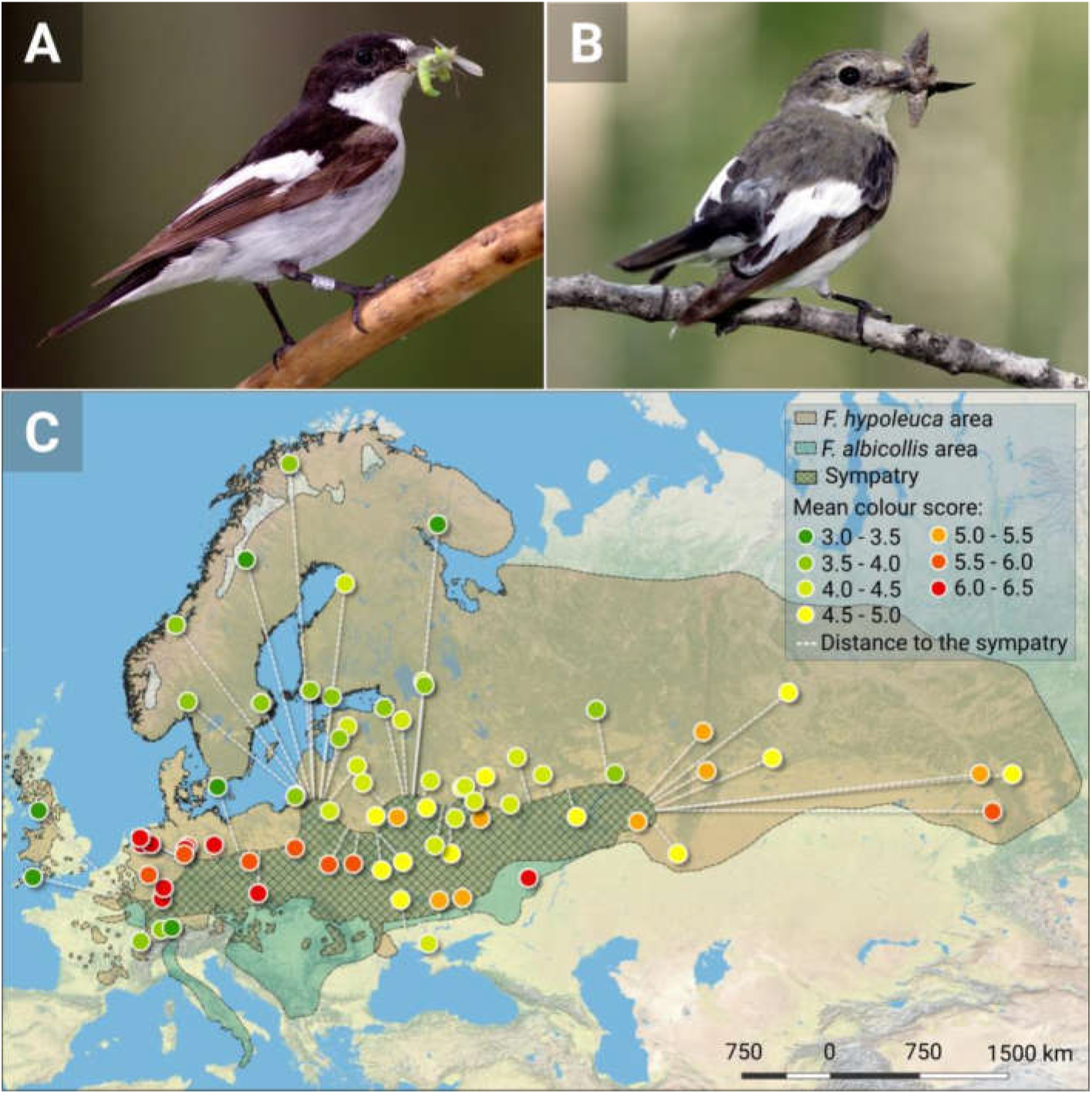
The variability of the male breeding plumage colour and the mean colour score in populations of the Pied Flycatcher (F. hypoleuca). (**A**) The contrast-coloured male (black morph). (**B**) The less bright male (brown morph). (**C**) The mean colour score in populations breeding in different parts of the Pied Flycatcher *F. hypoleuca* range. For each observation point, the distance to the border of the Collared Flycatcher *F. albicollis* is shown. Photos (**A**) and (**B**) credits: Vladimir G. Grinkov. Map source: the U.S. National Park Service (NPS) Natural Earth physical map.

In parallel with the publication of Sætre et al. (Saetre et al. 1997), it was nevertheless noted that the authors’ conclusions indeed are based on limited material. “Both the phylogeny and the data on mate choice are based on limited sampling. The case for reinforcement will be more secure when the generality of the preferences has been established, and when a more complete picture of the biogeographic history of the flycatchers is available”, wrote R. K. Butlin and T. Tregenza (Butlin and Tregenza 1997). However, these comments were not considered in the following studies (Noor 1999; Servedio and Noor 2003).

In this paper, we decided to follow the opinion of R. K. Butlin and T. Tregenza and conduct a biogeographic study of flycatchers on the largest material that we can obtain for analysis today. Specifically, we checked whether there is a relationship between the prevailing breeding plumage colour in males and the remoteness of the Pied Flycatcher population from the breeding area of the Collared Flycatcher. To do this, we collected all the data known to us about the phenotypic structure of all populations of the Pied Flycatcher currently studied. We managed to find information for 41 populations. In addition, we analysed zoological collections from 11 museums. This allowed us to further evaluate the phenotypic structure of another 38 metapopulations of the Pied Flycatcher. To check whether the distance from the nesting area of the Collared Flycatcher may have an impact on the phenotypic structure of the Pied Flycatcher, we used a random forest as spatial predictions framework (Hengl et al. 2018). The extensive data sampling and spatial analyses usage largely distinguish our work from similar attempts, in which limited sampling and the linear regression models were used for these purposes (Laaksonen et al. 2015).

## Material and Methods

### Basic methodological approach

We used point-referenced spatial data and a random forest for spatial predictions framework (RFsp) (Hengl et al. 2018) to find (1) the most important predictor variables highly related to the mean colour score of the males’ breeding plumage of the Pied Flycatcher populations (response variable), and (2) a good parsimonious prediction model of the response variable for any point of the breeding range of the Pied Flycatcher. The RFsp was chosen because it can obtain equally accurate and unbiased predictions as other methods of spatial analysis (generalized linear model, different versions of kriging, geographically weighted regression) (Čeh et al. 2018; Hengl et al. 2018). The RFsp is thought to be advantageous over the other spatial analyses for the purpose of this research, as the former needs no rigid statistical assumptions about the distribution and stationarity of the target variable, it is more flexible towards incorporating, combining and extending covariates of different types, and information overlap (multicollinearity) and over-parameterization is not a problem for RFsp (Hengl et al. 2018). The distances from “observation points” to the border of the breeding area of the Collared Flycatcher and geographical coordinates (latitude and longitude) of “observation points” are used as explanatory variables, thus incorporating geographical proximity and geographical connection effects into the prediction process (Hengl et al. 2018). The “observation points” are *F. hypolueca* population “centres” for which the mean breeding plumage colour score was obtained (see below). We used the average daily maximum temperature and its variability (standard deviation) as general climatic characteristics of the population habitat. We put thermal characteristics into the model as alternative explanatory variables, thus contrasting them with the distance to the sympatry area during the prediction process. We used thermal characteristics in the analysis because it was shown that there is a low air temperature depression of the advertising behaviour of brown males in the Pied Flycatcher (Ilyina and Ivankina 2001; Kerimov et al. 2014), and this can lead to an increase in the proportion of black males in the reproductive part of the Pied Flycatcher population in cold spring years (Kerimov et al. 2014). Additionally, the basal metabolic rate (the amount of energy per unit time that an individual must spend to keep the body functioning at rest) and fledgling production of the males’ morphs in the *F. hypoleuca* flycatcher were shown to be dependent on the ambient temperature (Sirkia et al. 2010; Kerimov et al. 2014).

### Data collection and preparation

#### Step I

As a source of temperature data, we used The National Centers for Environmental Prediction (NCEP) / National Center for Atmospheric Research (NCAR) Reanalysis data set R-1, and NCEP / Department of Energy (DOE) Reanalysis II data set R-2 (Kalnay et al. 1996; Kanamitsu et al. 2002). The NCEP/NCAR R-1 and NCEP/DOE R-2 are high-quality, well-documented, freely available state-of-the-art gridded reanalysis data sets with global coverage of many relevant atmospheric variables spanning 1957 to present and 1979 to present, respectively (Kemp et al. 2012). Data for many variables are available at 17 pressure levels ranging from 1000 to 10 mb. Other variables describe conditions either at or near the surface. These data have a spatial resolution of 2.5 x 2.5 degrees and a temporal resolution of 6 h (00, 06, 12, 18 h UTC). We used the **RNCEP** package (version 1.0.8) of functions (Kemp et al. 2012) in the open-source R language to access required temperature data. First, using the *NCEP.gather* function from the **RNCEP** package, we loaded surface temperature data (variable air.sig995 in data set) from April to May for each year in the interval spanning 1980 to 2015 for an area between 35- and 72-degrees north latitude and between −10- and 93-degrees longitude thus covering the breeding areas of the two species of flycatchers. Then using the *NCEP.restrict* function, we cut the interval from April 1 to April 14 and from May 16 to May 31, leaving the period from April 15 to May 15 for all loaded years. Finally, using the *NCEP.aggregate* function, we calculated the mean and standard deviation (SD) of daily maximum temperature for all loaded years, obtaining a point pattern of mean temperatures and SDs for the entire loaded area with a spatial resolution of 2.5 x 2.5 degrees of latitude x longitude. Similarly, we calculated the SD of the maximum daily temperature for May 1 for all years (the rendered thermal data map, see Fig. 2.). The latter assessment of variability mostly characterizes the repeatability or the predictability of temperature between years, and the former SD includes both inter-annual and intra-annual (inter-seasonal) variability, that is, a change in temperature from April 15 to May 15. Inter-seasonal temperature changes will be significantly higher for the continental climate than for the temperate climate. The selected interval spanning 1980 to 2015 seems to be sufficient both for obtaining an unbiased estimate of average temperatures and for a satisfactory assessment of its variability. The selected date interval, which lasts from April 15 to May 15, covers the period of birds’ arrival from African wintering grounds, pairing and the beginning of reproduction in most of the populations of considered species of flycatchers. Since the key principle of geography is that “everything is related to everything else, but near things are more related than distant things” (Miller 2004), the climatic characteristics of earlier or later periods or months (March or June, for example) will be highly correlated with the chosen one. Temporal and spatial autocorrelations of temperature data are well-founded and documented facts in modern science, for example, see (Di Cecco and Gouhier 2018). Our goal was not to understand the temperature of which the nesting period determines the phenotypic composition of the population to a greater degree (causation task); this is a task for other studies. We evaluated which variable better predicts the values of the response variable (correlation task). And in this sense, temporal and spatial autocorrelations of the temperatures are not a nuisance for us, but a feature that improves our approach, since it allows a rather arbitrary choice of the period for calculating thermal estimates.

**Fig. 2.**
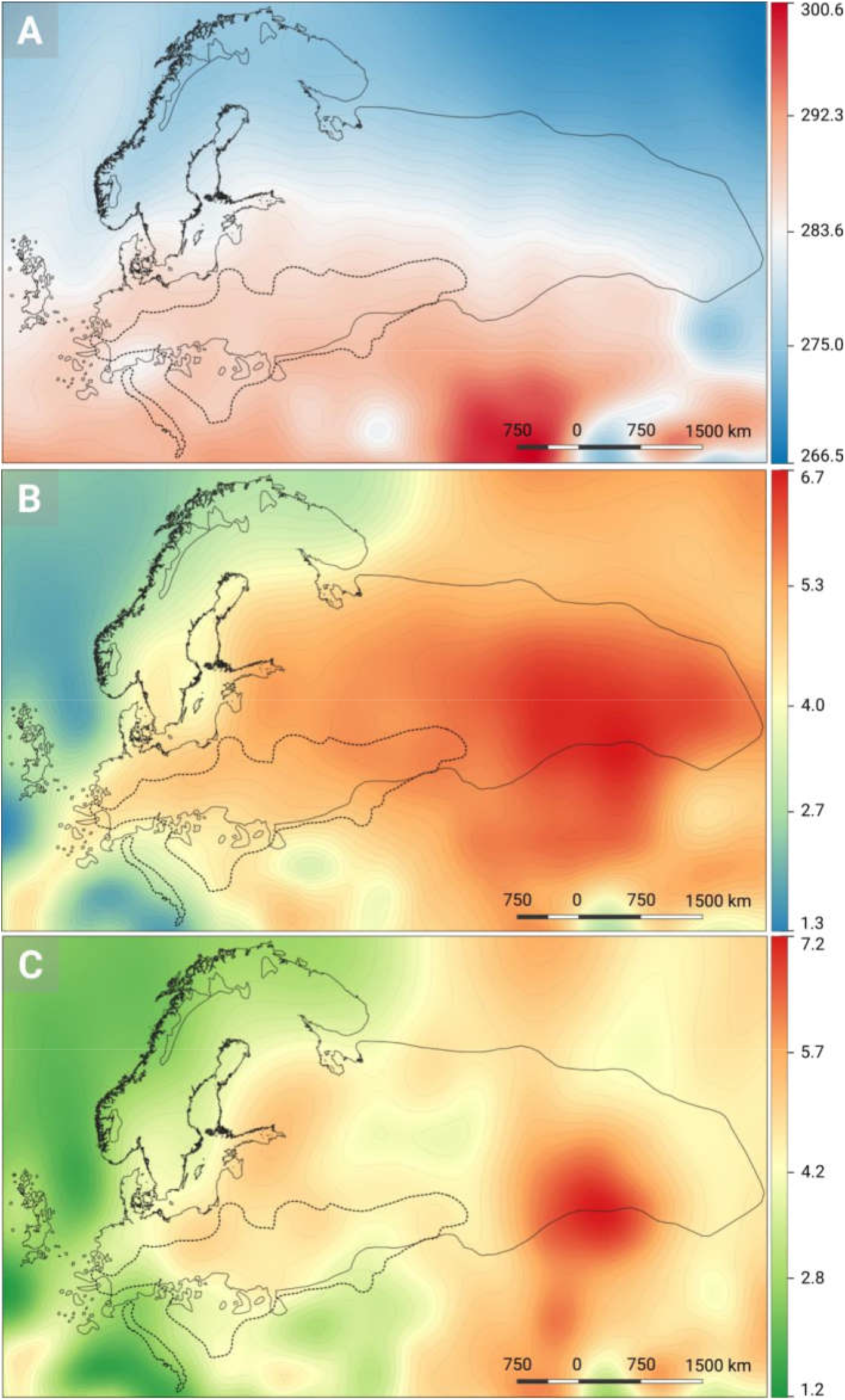
Thermal characteristics of the territories covered by the breeding ranges of the two species of *Ficedula* flycatchers. The images show (**A**) the mean daily maximum temperature in Kelvin, (**B**) the standard deviation of the daily maximum temperature, and (**C**) the standard deviation of the maximum temperature on May 1. The *F. hypoleuca* area is drawn with a solid line, the *F. albicollis* area is drawn with a dashed line. Each colour links to the value of the corresponding thermal characteristics (the key is on the right of each image). The hairlines connect the areas with the same values of the shown characteristic (for example, in the figure (**A**) these are isotherms).

#### Step II

We were looking for any articles studying the Pied Flycatcher that mentioned the phenotypic structure of this species. Seven variants (types or morphs) were initially identified in the variability range of the male breeding plumage colour in the Pied Flycatcher (Drost 1936). The value 1 of this scale was assigned to the bright coloured black and white males, and the value 7 assigned to the brown coloured males. Accordingly, it is easy to calculate the average colour score of the population, which will describe well the proportion of males of different morphs. The closer the average colour score is to 1, the greater number of black males there are, the blacker is the population as a whole and vice versa; the closer to 7, the greater number of brown males in the population, the browner is the population. In most of the publications, the mean colour score is given, therefore in our work we also used this indicator as the main descriptor of the phenotypic structure. We were able to find information about the phenotypic structure of the Pied Flycatcher for 41 populations (or metapopulations) (Table S1). For each, we found the nearest well-recognized geographical point. Usually it was nearest settlement to which the studied bird population was attributed by the author(s). We found the geographic coordinates in the QGIS program for this point. This geographic point was the “observation point” for those populations whose data on the phenotypic structure were obtained from publications. We used data from the same author (or the same group of authors) for one territory from only one most recent publication. If for the same territory (or a close region) we found data from different authors, then we used all such data in the calculations.

#### Step III

The Pied Flycatcher studies in the western part of the breeding range were carried out much more often than in the eastern part resulting in a shifted distribution of observation points (Table S1). To compensate for data bias, we examined the collections of the following museums: [1] Siberian Zoological Museum of the Institute of Systematics and Ecology of the Siberian Branch of the Russian Academy of Sciences (Novosibirsk), [2] Collection of the Department of Biogeography of the Faculty of Geography of the M. V. Lomonosov Moscow State University (Moscow), [3] State Museum of Nature of the Kharkiv National V.N. Karazin University (Kharkiv), [4] Zoological Museum of Kyiv National Taras Shevchenko University (Kiev), [5] Zoological Museum of Lviv National I. Frank University (Lviv), [6] Zoological Museum of the Belarusian State University (Minsk), [7] Zoological Museum of the M. V. Lomonosov Moscow State University (Moscow), [8] Zoological Museum of the National Academy of Sciences of Ukraine (Kiev), [9] Zoological Museum of the Zoological Institute of the Russian Academy of Sciences (St. Petersburg), [10] Kirov City Zoological Museum (Kirov), [11] Kaunas Zoological T. Ivanauskas Museum (Kaunas), [12] Museum of Natural History of Tartu University (Tartu).

We examined the bird skin collection and determined the colour type of bird skins in museums [1] – [9]. Data on the colour type of 17 males were kindly provided by V. N. Sotnikov from [10]. We received digital photographs of 12 and 11 bird skins from [11] and [12], respectively, for subsequent colour typing. In total, we examined 471 birds’ skins, of which 39 skins were excluded because some of them were clearly taken from migratory birds (autumn migration), and for some of them we could not accurately determine the location. Finally, we got colour types for 432 bird skins from museums [2] – [12] (no skin from museum [1] was included in the final data set; see Data S1). For each skin, the geographic coordinate was determined in the QGIS, based on the description of the place where the birds were caught written on the label.

Since the spatial resolution of temperature data was 2.5 x 2.5 degrees of a geographic coordinate, we used polygons of that size to form “populations” (small area populations) (Fig. S4). The minimum sample for a population was 5 individuals, because this number is enough to calculate the mean deviating by less than 1 point of the colour score (a prespecified error bound) with a probability of 0.95 from the population total mean in a sample of 366 individuals with a standard deviation 1.13 (the population total mean sample size and the standard deviation were estimated in step II). If fewer individuals fell into some 2.5 x 2.5 degrees polygon, we included such individuals in the nearest polygons defined by the QGIS. Such individuals should not be located from the polygons in which they were included at a distance greater than 1.25 degrees (Fig. S4). After this procedure, there were still individuals that did not form populations or were not included in a population. In such remaining cases, we gradually increased the size of the buffer from the geographical position of the individual, while the areas thus obtained began to overlap. If overlapping areas included 5 or more individuals, then we also considered them as the population for which the mean colour score was calculated (large area populations) (Fig. S4). We were unable to include 14 individuals in any of the populations using any of the above methods (Fig. S4). The “observation point” for populations formed by the skins from museum collections was calculated as the mean coordinate of all population components in QGIS (i.e. the centre of mass, and not the centroid which is the geometric centre of the population) (Fig. S4). We thus obtained 39 populations or observation points (Table S2).

As mentioned, not only breeding, but also migratory birds can get into museum collections. Also, the skin of birds can be collected purposefully of the same type or colour. It is relatively easy to separate autumn migrants by the bird life calendar and the plumage colouring (before the autumn migration, birds moult completely, and the colour of the post breeding plumage is quite different from the colour of the breeding one). However, in our opinion, it is impossible to separate spring migrants from local breeders. Thus, this problem was ignored by us, because we were unable to find criteria and procedures for cleaning the sample that would not lead to significant data loss. We found signs of directional collection of black males in only one case (the small standard deviation, a difference in the mean breeding plumage score from neighbouring populations) (Table S2). For one area in the sample of 6 individuals, there were five males with a colour score 2 and one male with a colour score 3. Such a ratio of morphs is almost impossible to obtain with a random selection method even in dark populations. This observation point was excluded from the analysis (Table S2).

#### Step IV

For two species of *F. hypoleuca* and *F. albicollis* flycatchers, we downloaded the distribution data (digitized georeferenced maps in geodatabase file format for use in GIS mapping software compiled by BirdLife International and Handbook of the Birds of the World) from BirdLife International (2018) IUCN Red List for birds, http://www.birdlife.org (downloaded from the site on 09.09.2018 using QGIS). We revealed that the bird findings from the Kola Peninsula, and from the northernmost and easternmost parts of the *F. hypoleuca* breeding area in Western Siberia, do not fall into the BirdLife International reproductive range of *F. hypoleuca*. To adjust the range of *F. hypoleuca*, we georeferenced in QGIS the species breeding area from the publication of Sirkia et al. (Sirkia et al. 2015). The final breeding range of *F. hypoleuca* basically contained the BirdLife International data, but the north-western borders were extended to the north and east to the coast of the Kola Peninsula, to the north of Western Siberia coinciding with the border from Sirkia et al. (Sirkia et al. 2015) for this region, and to the very east of Western Siberia thus including data from museum collections for the area.

We did not include the Spanish (Iberian) and north-western African Atlas Flycatcher in the range of *F. hypoleuca* (Fig. 1). The birds of these territories are well isolated from other European *F. hypoleuca* populations (they can occur in one territory during a migration only), and they are phenotypically closer to each other (Corso et al. 2015; Potti et al. 2016). The Iberian form seems to be intermediate between the African form and *F. hypoleuca* in morphology (Sangster et al. 2004). It is recommended that the Atlas Flycatcher *F. speculigera* should be separated from the Pied Flycatcher *F. hypoleuca* as a species, and the Iberian form should be considered as subspecies *F. hypoleuca iberiae* of *F. hypoleuca* (Sangster et al. 2004). Sometimes, Iberian birds are also distinguished as a separate species (Potti et al. 2016). In both *F. speculigera* and *F. h. iberiae*, the colour of the breeding plumage in males is mainly black-and-white, and its variability is much less than among birds from the rest of Europe (Corso et al. 2015).

We also corrected the north-eastern boundary of the *F. albicollis* breeding range (Moscow is not included in the nesting range of this species), using Vabishchevich and Formozov (Vabishchevich and Formozov 2010) georeferenced distribution data. Southern Sweden, the islands of Öland, Gotland and Saaremaa were also excluded from the final *F. albicollis* breeding range due to the recent expansion of the species over the Baltic Sea thus following Laaksonen et al. (Laaksonen et al. 2015) (Fig. 1).

#### Step V

All data obtained in the previous steps were entered QGIS for their final assembly. For each *F. hypoleuca* observation point, the distance in kilometres to the boundary of the *F. albicollis* range was calculated (Fig. 1). If the observation point fell inside the *F. albicollis* range, the distance got negative values. Binding of the *F. hypoleuca* observation point to the thermal characteristics of the territory was carried out using the method of the nearest neighbour (the nearest point-referenced values of the thermal characteristics of the territory were chosen), as well as by Inverse Distance Weighted (IDW) interpolation using the 4 nearest points. We used the GRASS 7 *v.surf.idw* algorithm for surface interpolation from point-referenced data by the IDW interpolation for which region the *cellsize* was set to 2.5 (according with the spatial resolution of the temperature data), the power parameter was set to 3, and the number of interpolation points was set to 4. Then we used the point sampling tool to assign the interpolated value to the population observation point.

Thus, our initial model consisted of one response variable (the mean breeding plumage colour of *F. hypoleuca* males in the population) and 9 predictors: *Y*-latitude, *X* - longitude, *cf.dist* - distance to the border of the *F. albicollis* range, *mt* - mean maximum daily temperature of the nearest temperature referenced point, *mt.idw* - IDW interpolated mean maximum daily temperature, *sd* - standard deviation of the maximum daily temperature from April 15 to May 15 spanning 1980 to 2015 of the nearest temperature referenced point, *sd.idw* - IDW interpolated standard deviation of the maximum daily temperature from April 15 to May 15 spanning 1980 to 2015, *sd.1d* - standard deviation of the maximum day temperature for May 1 spanning 1980 to 2015 of the nearest temperature referenced point, and finally *sd.1d.idw* - IDW interpolated standard deviation of the maximum day temperature for May 1 spanning 1980 to 2015.

### Spatial data analysis

We used the open-source R software environment for statistical computing and graphics (version 3.5.0) under an integrated development environment for R - RStudio (RStudio Desktop version 1.1.447) to analyse data assembled on step V. For regression tasks we used the **ranger** package (version 0.10.1) as an implementation of the random forests (Wright and Ziegler 2017). To obtain the most accurate predictions, the random forest parameters need to be optimised (Probst et al. 2018). To configure the parameters of the random forest, we used the **tuneRanger** (version 0.3) package (Probst et al. 2018) which allows model-based optimization for tuning strategy and the three parameters *min.node.size, sample.fraction* and *mtry* tuning at once. Out-of-bag predictions were used for evaluation. To evaluate the best *num.trees* parameter, we used the **caret** (version 6.0-80) package. We used a *trainControl* function from the package to set the resampling method to 70-fold cross-validation procedure. Validation was done by calculating the root mean squared error (RMSE). Thus, the initial model, which included all 9 predictors, was calculated with the following parameters of the random forest: *min.node.size=8, sample.fraction=0.79, mtry=2, num.trees=250, splitrule* = “variance”, importance=“impurity”, *num.threads=4, verbose=FALSE, respect.unordered.factors=TRUE*, replace=FALSE, *keep.inbag=TRUE*. We obtained the coefficient of determination for the whole model (R^2^) equal to 0.73 (out-of-bag predictions), and the mean squared error (MSE) equal to 0.24 (out-of-bag prediction error). To build the most parsimonious prediction model of the response variable, i.e. to reduce the number of predictors with minimal impact on the response variable, we conducted a nested cross-validation procedure using the **caret** package. We estimated the 10-fold cross-validated prediction performance of models with a sequentially reduced number of predictors ranked by variable importance (the variables with the lowest variable importance values was sequentially excluded from the models) in 100 repetitions (Fig. 3). In our case the variable importance was measured as the variance of the responses (Fig. 3C). Validation of the models was done by calculating the RMSE (Fig. 3A). The cross-validation procedure reduced 4 of 9 predictors (including the distance to the *F. albicollis* breeding area), and the final model performance increased slightly, but statistically significantly (Fig. 3A, B). Thus, the final model included only 5 predictors listed below: Y, X, *mt, sd, sd.1d*. After excluding 4 predictors from the model, we again conducted the above procedure for the parameter’s optimization of the random forest. The final model was calculated with the following parameters: *min.node.size=11, sample.fraction=0.787, mtry=2, num.trees=250, splitrule* = “variance”, importance=“impurity”, *num.threads=4*, verbose=FALSE, *respect.unordered.factors=TRUE*, replace=FALSE, *keep.inbag=TRUE*. The R^2^ for the whole final model is 0.75, and the MSE is 0.23. To predict the mean breeding plumage colour score of males in the *F. hypoleuca* within the entire breeding range (to calculate out of sample data), we used the *predict.ranger* function in the **ranger** package with the final random forest model. The values of all 5 predictors obtained in step 1 for the area between 35- and 72-degrees north latitude and between −10- and 93-degrees longitude were used. We calculated the standard error of the predictions using the infinitesimal jack-knife for bagging methodology (Wager et al. 2014) as it implemented in the **ranger** package (Fig. 4, Fig. S2).

**Fig. 3.**
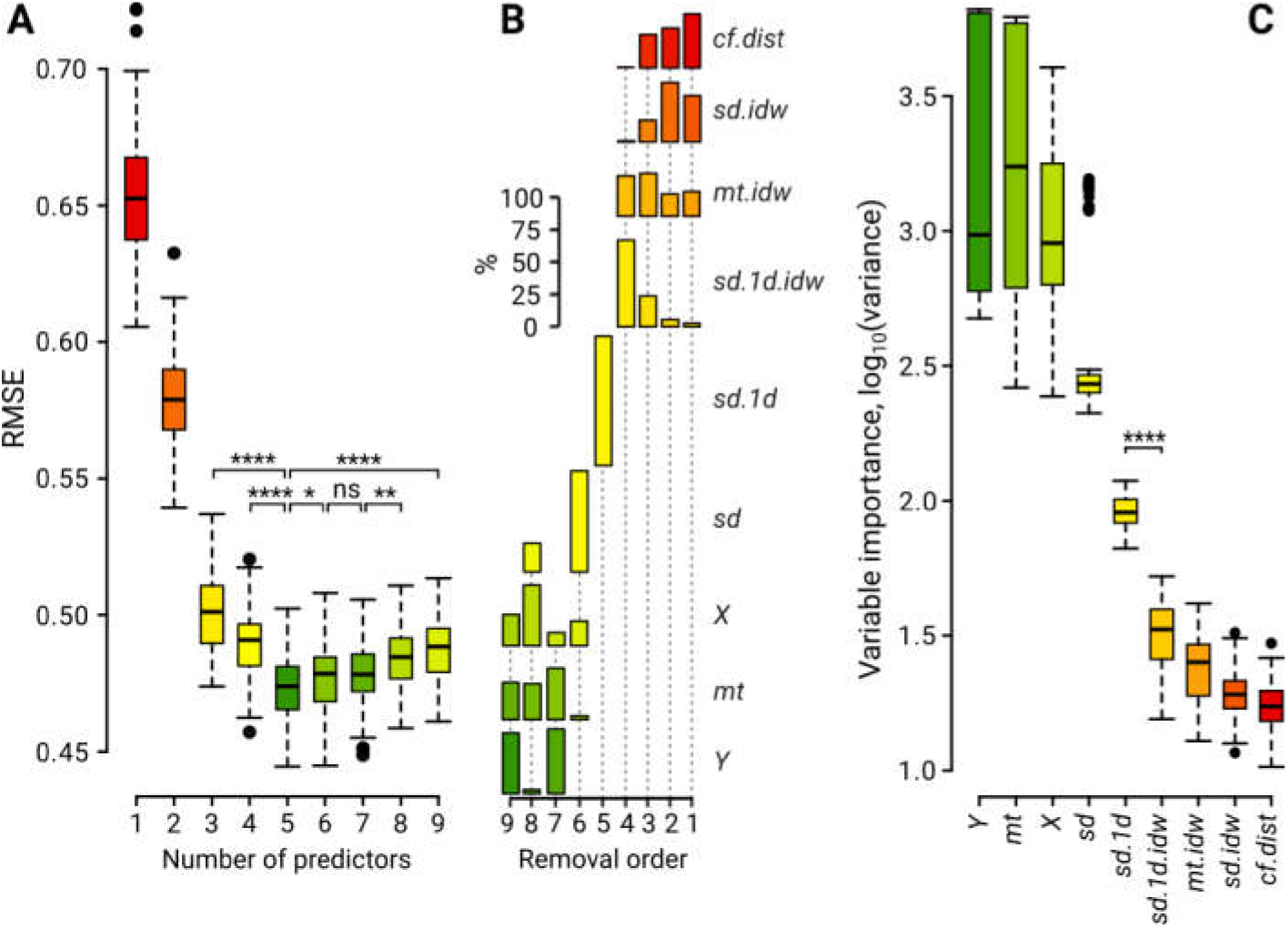
The results of the 10-fold cross-validation procedure for 100 repetitions. (**A**) Prediction performance of models calculated as the root mean squared error (RMSE) with sequentially reduced number of predictors. (**B**) The removal order of predictors and its frequency in percent (%), showing at what step and how often the predictors obtaining the lowest variable importance value were excluded from the models. (**C**) The variable importance values measured as the variance of the responses are given for all predictors (the variable importance values on the ordinate axis are converted by a decimal logarithm). The figure parts (**A**) and (**C**) show the median (the line across the box), the interquartile range (IQR, the box), the positive and negative 1.5*IQR extension of the IQR (the vertical dotted line). The small black circles represent outliers from the latter range. A horizontal line with downward serifs at the ends connects the compared values for which significance levels of Student’s t-test (two-tailed) shown as following: ns for p > 0.05; * for p < 0.05; ** for p < 0.01, and **** for p < 0.0001. Variable predictor names are in italics; for a description, please see the material and methods.

**Fig. 4.**
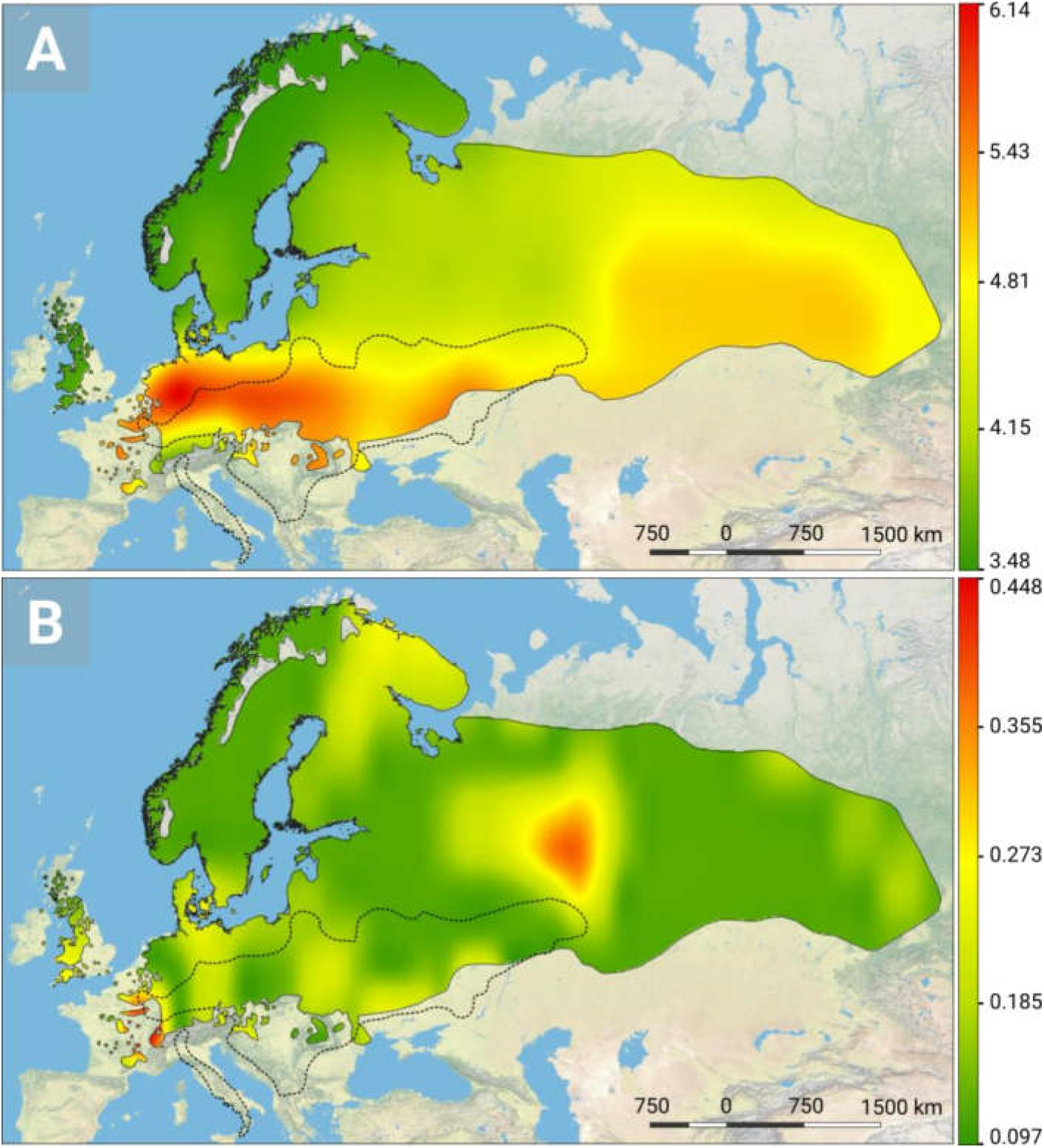
Reconstruction of the phenotypic structure of the *F. hypoleuca* populations for the entire breeding range. (**A**) Mean colour score predicted by the final random forest model. (**B**) Accuracy of the predicted values (the standard error of the predictions, see also Fig. S2). In the figure parts, the colour links to the value of the corresponding characteristic shown (the key is on the right of each part).

### Data visualization

To visualize the results of spatial analysis, we transformed point-referenced data into raster surface data in the QGIS. For this, we used the GRASS 7 *v.surf.idw* algorithm for surface interpolation from point-referenced data by the IDW interpolation (region *cellsize* was set to 2.5, power parameter was set to 3, and number of interpolation points was set to 4). To improve the spatial resolution of raster data and to improve the visualization effect, we used the SAGA Resampling algorithm with upscaling and downscaling IDW interpolation. We changed the *cellsize* parameter to obtain the most acceptable visualization effect (usually, the parameter value was equal or less than 0.1). To draw all the other graphs, we used the **ggplot2** (Wickham 2016) (version 3.0.0) package in R. Final processing of the vector-based drawings was carried out in Inkscape™: Open Source Scalable Vector Graphics Editor (version 0.92.3), and the raster-based drawings was done in GNU Image Manipulation Program (GIMP) (version 2.10.4).

### Modelling the influence of the *F. albicollis* flycatcher on the population phenotypic structure of the *F. hypoleuca* flycatcher

In the QGIS program, we modelled a hypothetical phenotypic structure of the *F. hypoleuca* populations, as if only one single factor was acting on the distance from the *F. albicollis* breeding range. This simulation was carried out only for visualization and was not used in statistical treatment. First, we transformed the *F. albicollis* range into a raster using the SAGA rasterize data/nodata algorithm, and then modelled the distance effect from the borders using the *r.grow.distance* algorithm with the cell size parameter set to 1. The final visualization was done using the above data visualization methods (Fig. S3).

### H. Löhrl data processing

The figure 2 given on page 271 of the H. Löhrl publication in Bonner Zoologische Beiträge (Löhrl 1965) was georeferenced in QGIS. We have downloaded freely accessible The Global Multi-resolution Terrain Elevation Data 2010 (GMTED2010) from the EarthExplorer site (https://earthexplorer.usgs.gov/) for the south-west of Germany and adjacent territories. We used the point sampling tool in QGIS to assign the terrain elevation values to the H. Löhrl observation points for further processing in R (Fig. 5).

**Fig. 5.**
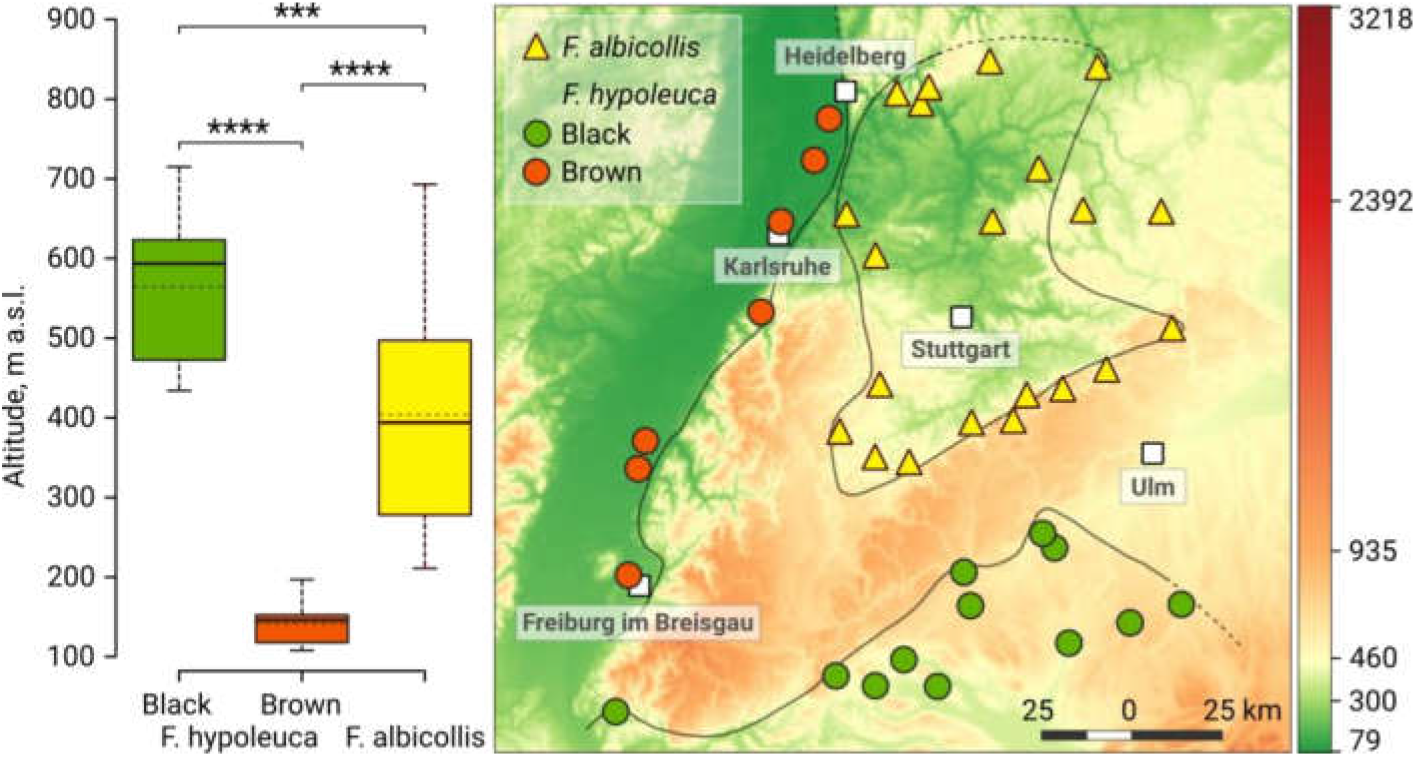
Small-scale spatial segregation between two species of the *Ficedula* flycatcher in south-west Germany, depending on the elevation above sea level. Reprocessed data of figure 2 (page 271) of the H. Löhrl publication in Bonner Zoologische Beiträge (Löhrl 1965) are shown. On the map, the colour indicates the height above sea level (altitude) in meters (according to the color rump on the right of the map) and the location of the nests of the two species of *Ficedula* flycatchers. The graph compares the height of the nest locations above sea level. The graph shows the median (the solid line across the box), the mean (the dashed line across the box), the interquartile range (IQR, the box), the positive and negative 1.5*IQR extension of the IQR (the vertical dotted line). A horizontal line with downward serifs at the ends connects the compared values for which the significance levels of Student’s t-test (two-tailed) are shown as following: *** for p < 0.001, and **** for p < 0.0001.

## Results

A simple graphical analysis of the relationship between the mean colour score and the predictors showed that there are obvious non-linear dependencies in the data structure, including the dependence of the response variable on the distance from the observation point to the sympatry area (Fig. S1). In this regard, the use of linear models cannot be considered as a quite adequate method of analysing such data (Laaksonen et al. 2015).

The thermal characteristics of the area covering the ranges of the flycatcher species demonstrate a strong, but quite expected pattern of variability. Within the Pied Flycatcher distribution area, there is a pronounced south-north gradient of average daily maximum temperatures (Fig. 2). It is particularly pronounced between Central and Eastern Europe on the one hand and Fennoscandia on the other. The variability and predictability of thermal characteristics has a more complex geographic pattern, although in general there is a tendency to increase unpredictability eastward (an increase in the climate continentality as one moves deeper into Eurasia) (Fig. 2).

We used a spatially explicit approach to achieve two main objectives. Firstly, for the purposes of interpretation, we found the most important variables highly related to the population mean colour score of the males’ breeding plumage of the Pied Flycatcher (response variable). Secondly, we found the smallest number of variables enough for a good parsimonious prediction of the mean colour score of the males’ breeding plumage for any point of the breeding range of the Pied Flycatcher.

The random forest regression has shown that the distance to the breeding area of the Collared Flycatcher is of minimal importance for predicting the phenotypic structure of the Pied Flycatcher populations (Fig. 3). The average daily maximum temperature during the breeding season and its variability are associated with the predominant male colour morph in each population of the species (Fig. 3). Latitude and longitude also affect the phenotypic structure of the Pied Flycatcher populations (Fig. 3). This may indicate that we did not include any other important variables associated with the geographical location of the population in the study.

Model predictions show that the predominance of black morphs in males is observed in the Pied Flycatcher populations nesting in areas with a predictably colder environment (Fennoscandia and the British Isles) (Fig. 4, Fig. S2). Brown-coloured males predominate in populations nesting in a relatively warm and/or very unstable climate. Populations with a predominance of brown morphs form the entire southern region of the breeding range of the Pied Flycatcher, including the most Eastern European part and the Western Siberian part. Moreover, the populations in which the average values of the colour score reach maximum values (more brown morphs in the population) are outside the breeding range of the Collared Flycatcher – these are central and northern Germany, and the Netherlands (Fig. 4). In sympatry, the phenotypic structure of the Pied Flycatcher populations also changes (Fig. 4), and these changes have a much more complex spatial pattern compared to what one would assume if the mere presence/absence of another species would influence the phenotypic structure of sympatric populations (Fig. S3).

## Discussion

The study of the mechanisms of speciation in the Old World flycatchers of the genus *Ficedula*, unfortunately, recalls the very case when the cart was putted before the horse. Acceptance of reinforcement as the main factor of speciation in the *Ficedula* flycatchers and as the mechanism of maintaining intraspecific variability in one of them (Saetre and Saether 2010) occurred even though the biogeographic pattern of phenotypic structure was not revealed for the whole breeding area of the interacted species, and the generality of the female preferences for male breeding plumage was not established.

Surprisingly, data on the mainly brown coloration of the males’ breeding plumage of the Pied Flycatcher in Western Siberia were available in the scientific literature (Johansen 1954). Apparently, this information was not widely known. This fact alone would be enough to cast doubt on the generality of the coupling between the character displacement and the areas of sympatry. Now we can say with high certainty that no convincing evidence of the coincidence of the character displacement and the sympatry areas of the flycatchers exist (Fig. 4). Our studies persuasively show that the geographic pattern of variability of the mean colour score of males’ breeding plumage in the Pied Flycatcher is much more complicated and cannot be caused solely by interspecific interactions (Fig. 1, Fig. 4 and Fig. S3). For example, the shift of male breeding plumage colouration occurs in the sympatry, but also in the allopatry in most eastern parts of the breeding range of the Pied Flycatcher up to the eastern distribution boundaries in Western Siberia, that is, the most remote populations from the areas of sympatry (Fig. 1 and Fig. 4).

It is crucially important that if reinforcement is inferred from displacement in a secondary male sexual character, the displacement must be detectable by females; and females should use these characters for a mate choice (Howard 1993; Noor 1999). In addition to the publication of Sætre et al. (Saetre et al. 1997), some other studies reveal that a female preferably chooses a black male in allopatric populations of the Pied Flycatcher (Roskaft and Jarvi 1983; Saetre et al. 1994; Dale and Slagsvold 1996). However, there are also several studies done by independent researchers about different allopatric populations in which no clear benefits for black morphs in mate choice were found (Alatalo et al. 1986; Alatalo et al. 1990b; Potti and Montalvo 1991; Lehtonen et al. 2009; Sirkia and Laaksonen 2009). This indicates that the generality of the female preferences for male breeding plumage in the Pied Flycatcher also cannot be considered a fully proven phenomenon.

We think that if the female perceives the male breeding plumage colour as a cue to select a mate, this male trait is not the main one for mate choice or not used at all. For example, it was recently shown that the intensity of the advertising behaviour of free-living males of the Pied Flycatcher is modified by ambient air temperatures in the Moscow Region population (Russia, European part). Black males can maintain a high intensity of advertising behaviour at both low and high temperatures. However, brown males are able to attract females effectively only at higher air temperatures, and at low temperatures reduce dramatically the intensity of advertising behaviour (Ilyina and Ivankina 2001; Kerimov et al. 2014). Quite similar results were obtained in the experiments in the Tomsk population (Russia, Western Siberia) of the Pied Flycatcher. It was shown that females choose black males at low ambient temperatures merely because they are more active (Kerimov et al. 2014). Moreover, it was possible to demonstrate that this mechanism influences the morph ratio in the reproductive part of the population – in years with a cold spring, more black males enter the reproductive part of the Tomsk population, apparently due to the temperature-related depression of the brown males’ advertising behaviour (Kerimov et al. 2014). In the literature, there is also evidence that the ambient temperature can affect the reproductive success and the energetics of the morphs. The reproductive output of black males was shown to be the highest when it was cold during egg-laying but warm during the nestling period, whereas the fledgling production of brown males was highest when it was continuously warm (Sirkia et al. 2010). Brown males increase a basal metabolic rate (BMR; the amount of energy per unit time that an individual must spend to keep the body functioning at rest) under the influence of low ambient temperatures, and black males, on the contrary, retain the same level of BMR in a very wide range of daily temperatures (Kerimov et al. 2014). It is likely that the opposed results of earlier studies of female morph preferences could be obtained because the temperature dependence of the males’ energetics and behaviour was not considered. To date, data from Sætre et al. (Saetre et al. 1997) remain the only “strong” evidence of reinforcement (Saetre and Saether 2010) (all others are not an exclusive prerogative of reinforcement) (Noor 1999; Butlin and Smadja 2018). Therefore, the replication of the experiments of Sætre et al. (Saetre et al. 1997) to check the female preferences is necessary on a more extensive material, considering the new information about the dependence of the males’ advertising behaviour on ambient temperatures.

The coupling of males’ breeding plumage colour variability with weather characteristics in the Pied Flycatcher was noted in one of the first studies devoted to this issue. Löhrl (Löhrl 1965) described a very interesting picture of the spatial segregation of male morphs of the Pied Flycatcher and the Collared Flycatcher. He showed that in southwest Germany brown and black males of the Pied Flycatcher are separated from each other spatially, as well as from the Collared Flycatcher (Fig. 5). Brown males occupied the forests of the lowest lands, the Collared Flycatcher occupied the forests at medium height above sea level, and the black males nested in the highest mountain forests. Such a clear spatial segregation between two species and morphs cannot be explained by a simple interference competition over nest cavities, where the collared flycatcher is a winner (Saetre and Saether 2010). This habitat segregation on such a spatial scale can only be the result of an active habitat choice. Indeed, there is experimental evidence for species-specific habitat preferences in two flycatcher species in their hybrid zone (Adamik and Bures 2007).

All these data can be regarded as evidence considering that the evolution of the plumage colour in the Pied Flycatcher is not driven by reinforcement but is the result of the adaptation of black males to the conditions of higher mountains forests, and brown ones – to the conditions of lower-altitude forests. Then the distribution of brown and black populations of the Pied Flycatcher along the modern breeding area is a projection of adaptations to different ecological subniches that the species have developed in refugia during the last glaciation (*F. albicollis* in Italy, and *F. hypolueca* on the Iberian Peninsula) and their subsequent evolution (Saetre and Saether 2010).

It is very likely that the initial stages of the *F. hypolueca* species formation took place in the relatively higher-altitude conditions of the north of the Iberian Peninsula refuge during Quaternary ice ages (for the European paleoenvironments and maximum extent of ice, for example, see (Tzedakis et al. 2013). Further, as the glaciation receded, the *F. hypolueca* began to expand earlier along its current range, since there are no physical obstacles in this refugium like the Alps in the north of the Apennine Peninsula. Probably, as a result of this expansion, an adaptation to breeding in lower-altitude forests evolved, marked by the brown colour of the male breeding plumage. The expansion of the *F. albicollis* from its Italian refuge could be limited to the Alps for some time. As a result, when *F. albicollis* began to spread over its existing range, it encountered the brown *F. hypolueca* populations that had spread a little bit earlier. In contact zones the species effectively diverged to different ecological niches, although the *F. albicollis* remained a more stenobiont species (Qvarnstrom et al. 2010). At present, the species are so well segregated ecologically that there may be no competition for microhabitats and nesting sites between them (Walankiewicz et al. 1997; Adamik and Bures 2007; Czeszczewik et al. 2012). Therefore, it seems very likely that the black *F. hypolueca* populations have never shared the same habitats with the *F. albicollis* throughout the entire evolutionary history of these species! Thus, at present the British Isles and Fennoscandia are inhabited by black males preadapted to higher-altitude forests (ancestral state), and the southern and eastern parts of the Pied Flycatcher range are inhabited by brown males adapted to lower-altitude forests (newly evolved state). Thus, there is a clear ecological background for the evolution of the breeding plumage colour in the Pied Flycatcher males as an alternative for reinforcement including ecological adaptation, niche differentiation, and ecological character displacement.

In theoretical studies it was noted that in order to attribute mate choice patterns to reinforcement it was necessary that the character displacement must not have occurred for other, especially ecological reasons (Howard 1993; Noor 1999). This requirement is one where many studies have failed (Howard 1993; Noor 1999), and the reinforcement in *Ficedula* flycatchers seems to have to fill their numbers.

## Acknowledgments

We are very grateful to our colleagues from museums and universities for their support in collecting data: A.M. Peklo, L.N. Prokopchuk, A.A. Atemasov, V.F. Chernikov, Y.A. Redkin, P.S. Tomkovich, M.V. Kalyakin, L.G. Emelyanova, A.D. Pisanenko, E.A. Srebrodolskaya, A.V. Bokotey, V.M. Loskot, V.N. Sotnikov, A.I. Milutin, S. Rumbutis, A.P. Vabishchevich. We are thankful to N.A. Formozov for consultations. We are grateful to K. Henne for improving the English language of an earlier version of the manuscript.

## Funding

This work was supported by the Russian Fund of Basic Research RFBR (project 18-04-00536-a) and the State Assignment Ch. 2 CITIS AAAA-A16-116021660031-5.

## Author contributions

The conceptualization, data curation, funding acquisition, investigation, methodology, resources, validation, visualization, review, and editing were equally distributed among co-authors. Besides the listed, V.G. Grinkov was responsible for supervision, project administration, formal statistical analysis, and writing the original draft.

## Competing interests

The authors declare no competing interests.

## Data and materials availability

All data and software are available in the main text, the supplementary materials, and on the internet sites for purposes of reproducing the results or extending the analysis.

## Software and data sources

Quantum GIS Development Team. 2013. QGIS Geographic Information System. Open Source Geospatial Foundation Project. https://qgis.org

R Core Team. 2018. R: A language and environment for statistical computing. R Foundation for Statistical Computing, Vienna, Austria. https://www.r-project.org/

RStudio Team. 2016. RStudio: Integrated Development for R. RStudio, Inc. Boston, MA. http://www.rstudio.com/.

Environmental Prediction (NCEP)/National Center for Atmospheric Research (NCAR) Reanalysis data set R-1. URL http://www.cpc.ncep.noaa.gov/products/wesley/reanalysis.html

Environmental Prediction (NCEP)/Department of Energy (DOE) Reanalysis II data set R-2. URL http://www.cpc.ncep.noaa.gov/products/wesley/reanalysis2/index.html

Global Multi-resolution Terrain Elevation Data 2010 (GMTED2010). the EarthExplorer. URL https://earthexplorer.usgs.gov/

Species distribution data compiled by BirdLife International and Handbook of the Birds of the World from BirdLife International (2018) IUCN Red List for birds. URL http://www.birdlife.org

## Supplementary Materials

**Fig. S1.**
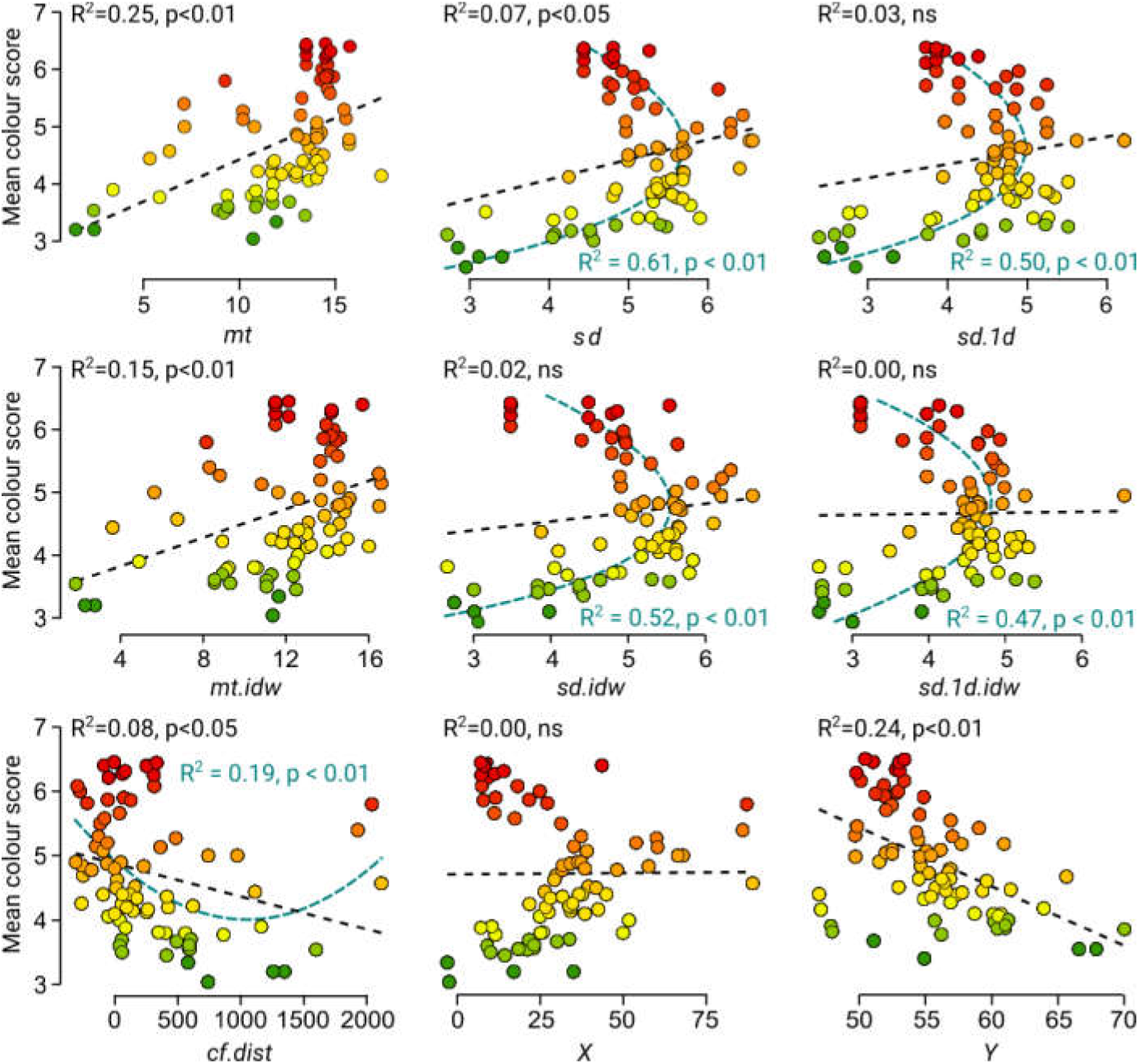
The relationship between the mean colour score (response variable) and all predictors. The scatterplots are presented for each predictor separately (the predictor name is written in italics under the x-axis), the linear (black dashed line) and non-linear (blue dashed curve) regressions are drawn; for each regression, the coefficient of determination (R^2^) and the significance level are given (indicated in black and blue font, respectively).

**Fig. S2.**
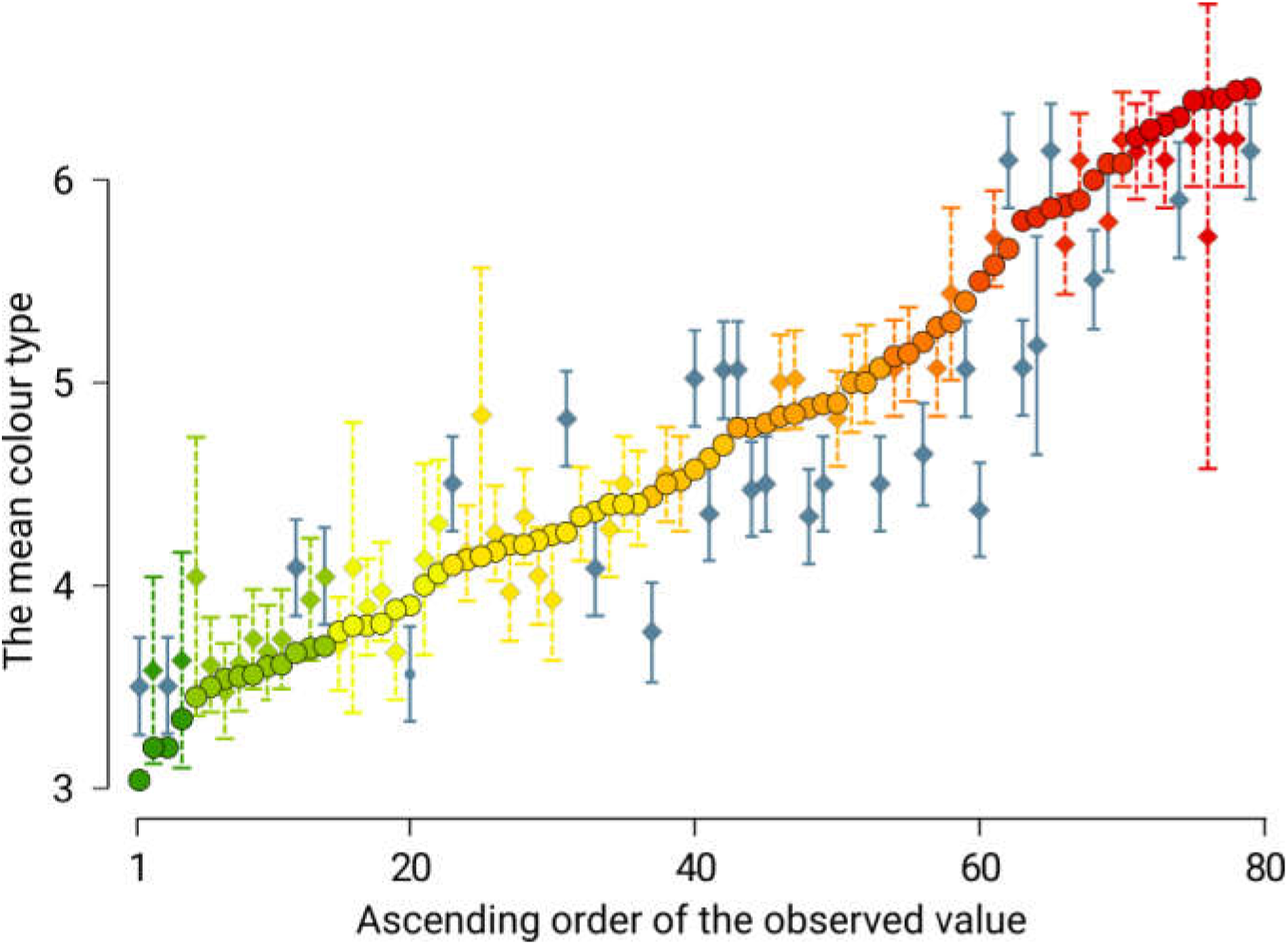
Comparison of observed and predicted values of mean colour score of male breeding plumage in the *F. hypoleuca* populations. The observed values are drawn in circles, the corresponding predicted values are drawn in diamonds; for each predicted value, a dashed vertical line indicates a 95% confidence interval (95% CI). Dark blue colour indicates such predictions that do not contain the observed value in the 95% CI.

**Fig. S3.**
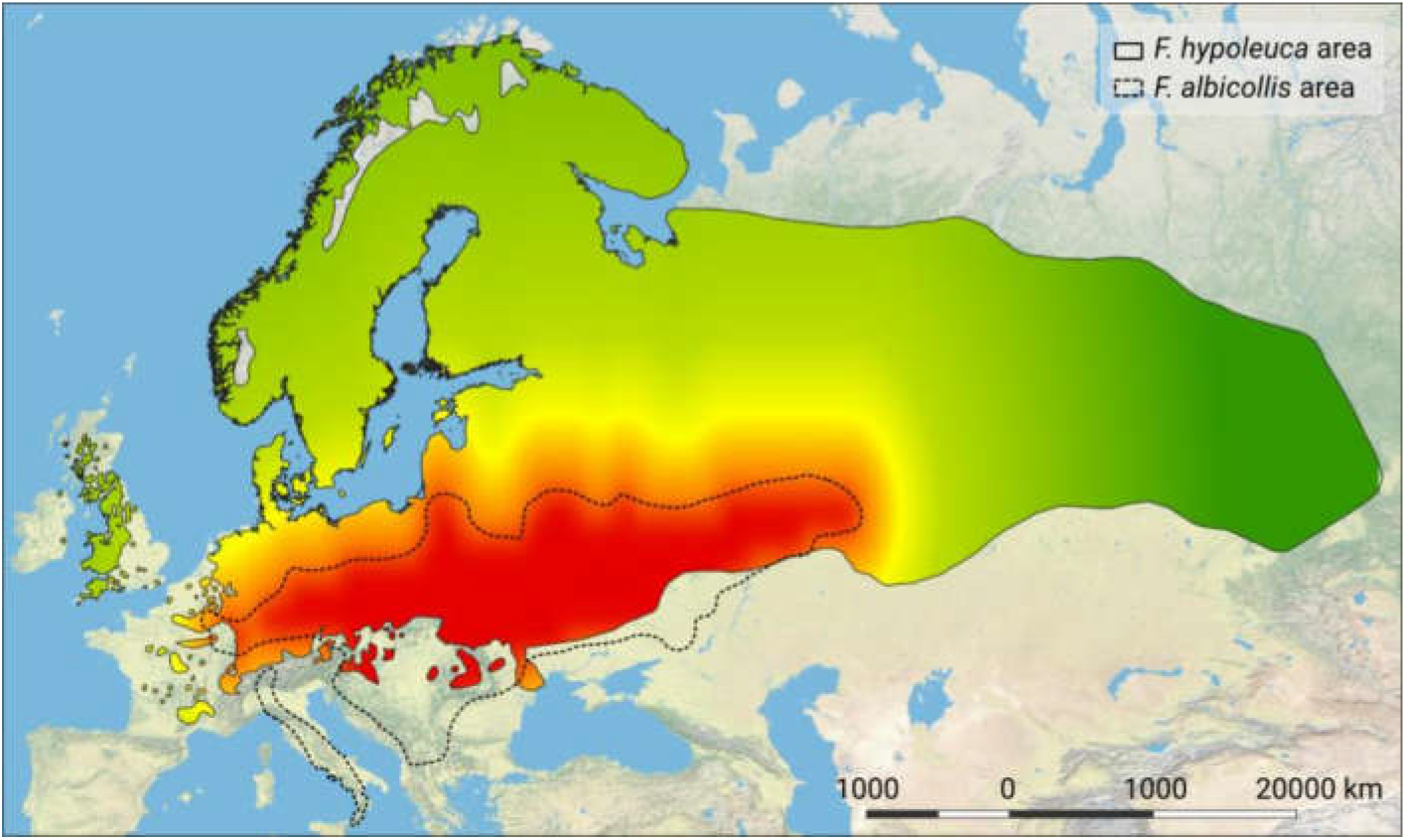
The hypothetical phenotypic structure of *F. hypoleuca* populations is drawn as if it was determined by the distance from the nesting range of *F. albicollis*. The transition from red through yellow to green corresponds to a change in the proportions of different morphs in the populations of *F. hypoleuca* from the most frequent brown through the intermediate to the most frequent black, respectively.

**Fig. S4.**
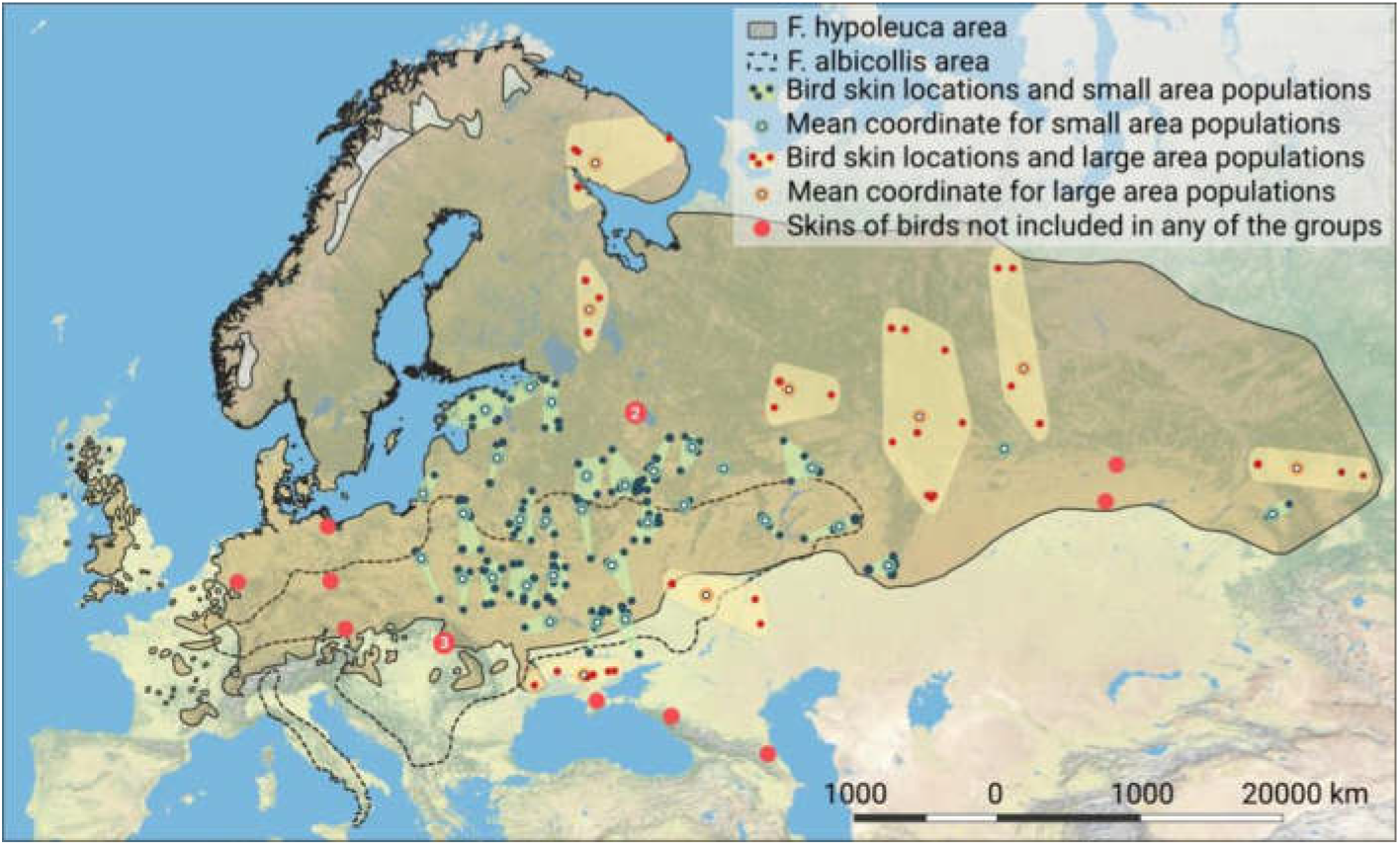
“Populations” formed from bird skins which were studied in 11 museums. The average population coordinate is the observation point used in the calculations (for summary data, please, see Table S2).

**Table S1.** The mean colour of males’ breeding plumage in the Pied Flycatcher *Ficedula hypoleuca* populations.

| # | Longitude | Latitude | Mean colour type | Sample size | Location | Country | Source |
| --- | --- | --- | --- | --- | --- | --- | --- |
| 1 | 17.36 | 49.64 | 6.08 | 50 | Moravia-Jeseníky | Czech Republic | Laaksonen et al. 2015; Sirkia et al. 2015 |
| 2 | 24.95 | 58.15 | 3.81 | 73 | Kilingi-Nõmme | Estonia | Laaksonen et al. 2015; Sirkia et al. 2015 |
| 3 | 22.16 | 60.43 | 3.56 | 408 | Ruissalo | Finland | Laaksonen et al. 2015; Sirkia et al. 2015 |
| 4 | 22.24 | 60.44 | 3.61 | 1207 | Turku | Finland | von Haartman 1985 |
| 5 | 24.25 | 60.14 | 3.80 | 117 | Siuntio | Finland | Roskaft et al. 1986 |
| 6 | 25.50 | 65.02 | 4.44 | 48 | Oulu | Finland | Laaksonen et al. 2015 |
| 7 | 7.09 | 50.74 | 5.86 | 66 | Bonn | Germany | Ristow 1975 |
| 8 | 7.29 | 52.52 | 6.40 | 416 | Lingen | Germany | Winkel and Winkel 1984 |
| 9 | 7.32 | 52.53 | 6.39 | 806 | Lingen | Germany | Handbuch der Vögel Mitteleuropas, 1993 |
| 10 | 8.36 | 49.30 | 6.21 | 42 | Harthausen | Germany | Laaksonen et al. 2015; Sirkia et al. 2015 |
| 11 | 8.51 | 49.96 | 6.45 | 49 | Mönchbruch, Hessen | Germany | Trettau 1952/53 |
| 12 | 10.43 | 51.94 | 5.66 | 44 | Goslar, Harz, Lower Saxony | Germany | Zang 1975 |
| 13 | 10.52 | 52.25 | 6.27 | 536 | Braunschweig | Germany | Winkel et al. 1970 |
| 14 | 10.78 | 52.42 | 5.90 | 1583 | Wolfsburg, Lower Saxony | Germany | Sternberg et al. 2002 |
| 15 | 13.24 | 52.49 | 6.31 | 15 | Berlin, Grunewald | Germany | Curio 1959 |
| 16 | 27.15 | 55.89 | 4.34 | 60 | Kraslava | Latvia | Laaksonen et al. 2015; Sirkia et al. 2015 |
| 17 | 9.65 | 63.33 | 3.90 | 166 | Songli | Norway | Roskaft et al. 1986 |
| 18 | 10.75 | 59.92 | 3.77 | 72 | Oslo | Norway | Laaksonen et al. 2015; Sirkia et al. 2015 |
| 19 | 20.29 | 69.36 | 3.54 | 95 | Skibotn | Norway | Laaksonen et al. 2015; Sirkia et al. 2015 |
| 20 | 16.55 | 51.52 | 5.58 | 105 | Jemielno, Silesia | Poland | Trettau and Merkel 1943 |
| 21 | 20.83 | 55.16 | 3.69 | 459 | Courish spit | Russia | Vysotsky 1995 |
| 22 | 32.80 | 60.85 | 4.22 | 217 | Scientific field station "Mayachino" | Russia | Laaksonen et al. 2015; Sirkia et al. 2015 |
| 23 | 32.94 | 60.68 | 3.70 | 93 | Ladoga Ornithological Station "Gumbaricy" | Russia | Grinkov 1997 |
| 24 | 33.89 | 52.45 | 4.26 | 110 | Bryansky Les Nature Reserve | Russia | Own data |
| 25 | 35.81 | 53.98 | 4.40 | NA | Ugra National Park | Russia | Vabishchevich and Formozov 2010 |
| 26 | 36.72 | 55.70 | 4.40 | 1403 | Zvenigorod | Russia | Ivankina et al. 2007 |
| 27 | 37.57 | 54.90 | 4.80 | 1631 | Moscow region | Russia | Anorova 1977 |
| 28 | 37.60 | 54.90 | 4.10 | NA | PTS Natural Biosphere Reserve | Russia | Vabishchevich and Formozov 2010 |
| 29 | 59.34 | 56.49 | 5.13 | 376 | Ekaterinburg | Russia | Belskii and Lyakhov 2004 |
| 30 | 84.95 | 56.35 | 5.40 | 1770 | Tomsk | Russia | Own data |
| 31 | 13.55 | 55.67 | 3.45 | 54 | Vombs fure | Sweden | Laaksonen et al. 2015 |
| 32 | 16.21 | 65.95 | 3.20 | 403 | Ammarnäs | Sweden | Nyholm and Myhrberg 1983 |
| 33 | 17.63 | 59.85 | 3.55 | 408 | Uppsala | Sweden | Lundberg and Alatalo 1992 |
| 34 | 6.35 | 46.63 | 3.88 | 81 | Vaud | Switzerland | Laaksonen et al. 2015; Sirkia et al. 2015 |
| 35 | 8.29 | 47.39 | 3.60 | 34 | Aargau-Zürich | Switzerland | Sternberg 1964 |
| 36 | 9.27 | 47.49 | 3.50 | 73 | Bischofszell, Thurgau | Switzerland | Eggenberger 1964 |
| 37 | 6.31 | 52.88 | 6.44 | 112 | Diever - Doldersum | The Netherlands | Bijlsma and Both 2014 |
| 38 | 6.42 | 52.52 | 6.25 | 77 | Ommen | The Netherlands | Haverschmidt 1973 |
| 39 | 6.60 | 52.86 | 6.08 | 440 | Drenthe | The Netherlands | Laaksonen et al. 2015; Sirkia et al. 2015 |
| 40 | -3.73 | 50.60 | 3.34 | 88 | East Dartmoor | United Kingdom | Laaksonen et al. 2015; Sirkia et al. 2015 |
| 41 | -3.20 | 54.40 | 3.04 | 489 | Cumbria | United Kingdom | Lundberg and Alatalo 1992 |

**Table S2.** The observation points for “populations” formed by the bird skins from museum collections. The mean colour type for population with ID 4, indicated in square brackets, was excluded from the analysis (see the materials and methods for the explanation). The column “Populations” contains identifiers of populations, denoted as L-1, L-2, etc. for large populations and S-1, S-2, etc. for small populations, respectively (please also see Fig. S4 and Data S1).

| ID | Longitude | Latitude | Mean colour type | Standard deviation | Number of the bird skins | Populations |
| --- | --- | --- | --- | --- | --- | --- |
| 1 | 33.29 | 46.49 | 4.14 | 1.46 | 7 | L-1 |
| 2 | 42.67 | 50.55 | 6.40 | 0.89 | 5 | L-2 |
| 3 | 34.17 | 67.26 | 3.20 | 1.10 | 5 | L-3 |
| 4 | 33.68 | 62.50 | [2.17] | 0.41 | 6 | L-4 |
| 5 | 49.00 | 59.54 | 3.80 | 1.10 | 5 | L-5 |
| 6 | 59.02 | 58.47 | 5.27 | 1.56 | 11 | L-6 |
| 7 | 67.00 | 60.35 | 5.00 | 1.26 | 6 | L-7 |
| 8 | 87.96 | 56.34 | 4.57 | 0.98 | 7 | L-8 |
| 9 | 20.81 | 52.30 | 5.59 | 1.28 | 17 | S-1 |
| 10 | 20.92 | 55.24 | 4.25 | 1.06 | 12 | S-2 |
| 11 | 23.97 | 51.35 | 6.00 | 1.20 | 8 | S-3 |
| 12 | 24.05 | 54.37 | 3.91 | 0.94 | 11 | S-4 |
| 13 | 26.25 | 51.40 | 5.82 | 1.47 | 11 | S-5 |
| 14 | 26.62 | 56.76 | 4.13 | 1.36 | 8 | S-6 |
| 15 | 25.73 | 58.73 | 4.20 | 1.23 | 10 | S-7 |
| 16 | 28.91 | 51.01 | 4.69 | 1.11 | 13 | S-8 |
| 17 | 28.38 | 54.10 | 4.63 | 1.60 | 8 | S-9 |
| 18 | 29.16 | 59.63 | 3.67 | 0.82 | 6 | S-10 |
| 19 | 30.69 | 49.23 | 4.78 | 1.20 | 9 | S-11 |
| 20 | 30.90 | 51.50 | 4.85 | 1.09 | 39 | S-12 |
| 21 | 30.39 | 54.04 | 5.50 | 0.93 | 8 | S-13 |
| 22 | 30.80 | 59.05 | 4.36 | 1.21 | 11 | S-14 |
| 23 | 34.30 | 49.21 | 5.14 | 1.10 | 14 | S-15 |
| 24 | 33.12 | 54.55 | 4.88 | 1.81 | 8 | S-16 |
| 25 | 33.48 | 56.00 | 4.20 | 1.79 | 5 | S-17 |
| 26 | 36.48 | 49.36 | 5.30 | 1.06 | 10 | S-18 |
| 27 | 35.41 | 51.98 | 4.90 | 1.79 | 10 | S-19 |
| 28 | 36.45 | 55.60 | 4.94 | 1.30 | 17 | S-20 |
| 29 | 38.17 | 53.95 | 5.07 | 1.49 | 14 | S-21 |
| 30 | 38.64 | 56.21 | 4.50 | 1.49 | 48 | S-22 |
| 31 | 40.95 | 54.73 | 4.50 | 1.38 | 6 | S-23 |
| 32 | 41.58 | 57.21 | 4.17 | 0.41 | 6 | S-24 |
| 33 | 44.00 | 56.32 | 4.40 | 1.34 | 5 | S-25 |
| 34 | 47.18 | 54.06 | 4.78 | 1.72 | 9 | S-26 |
| 35 | 52.89 | 53.79 | 5.25 | 1.28 | 8 | S-27 |
| 36 | 50.76 | 56.36 | 4.00 | 1.63 | 13 | S-28 |
| 37 | 56.64 | 51.97 | 4.83 | 1.40 | 12 | S-29 |
| 38 | 65.50 | 57.17 | 5.00 | 1.58 | 5 | S-30 |
| 39 | 86.11 | 54.34 | 5.80 | 1.30 | 5 | S-31 |

**Data S1.**
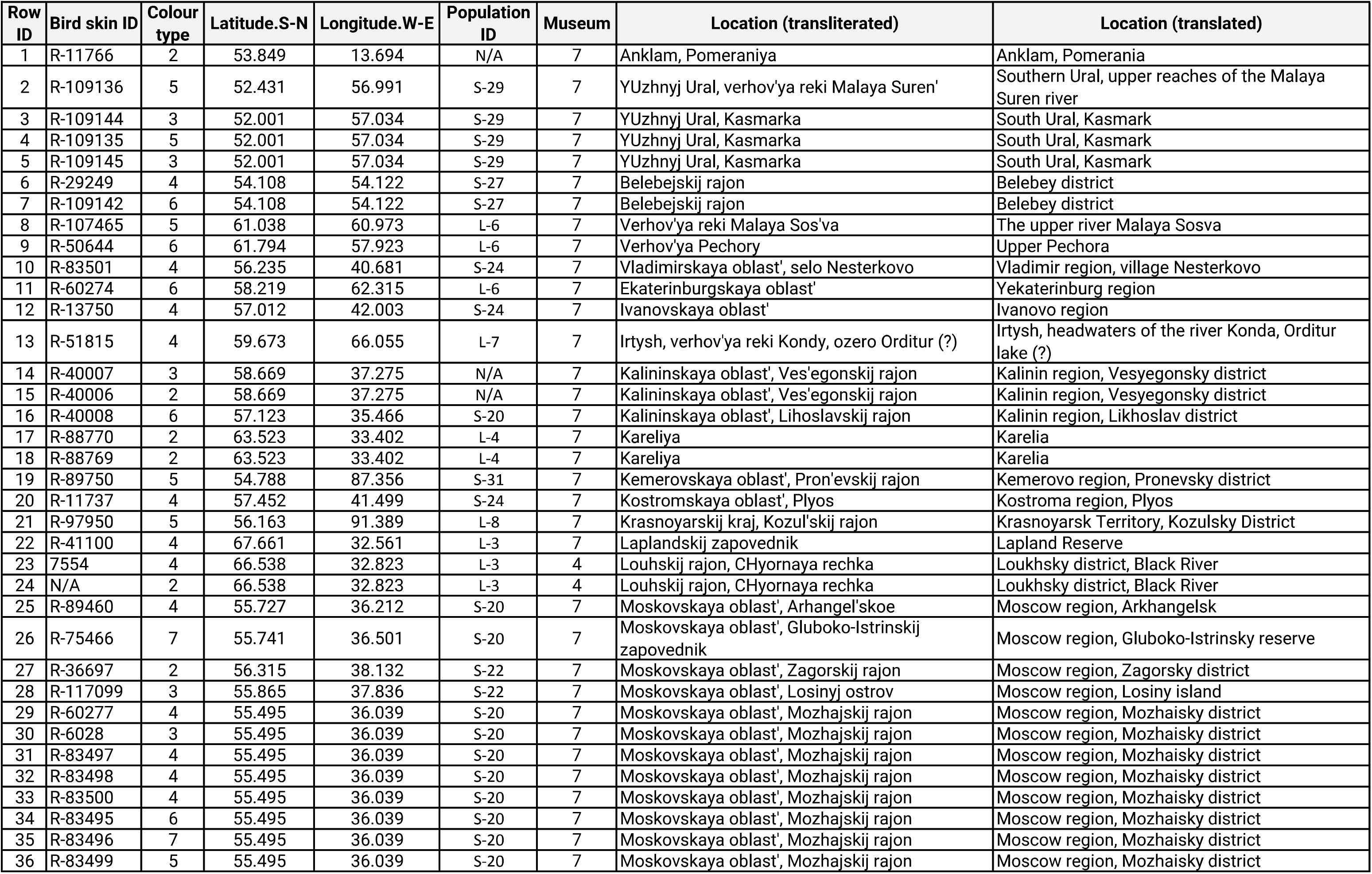
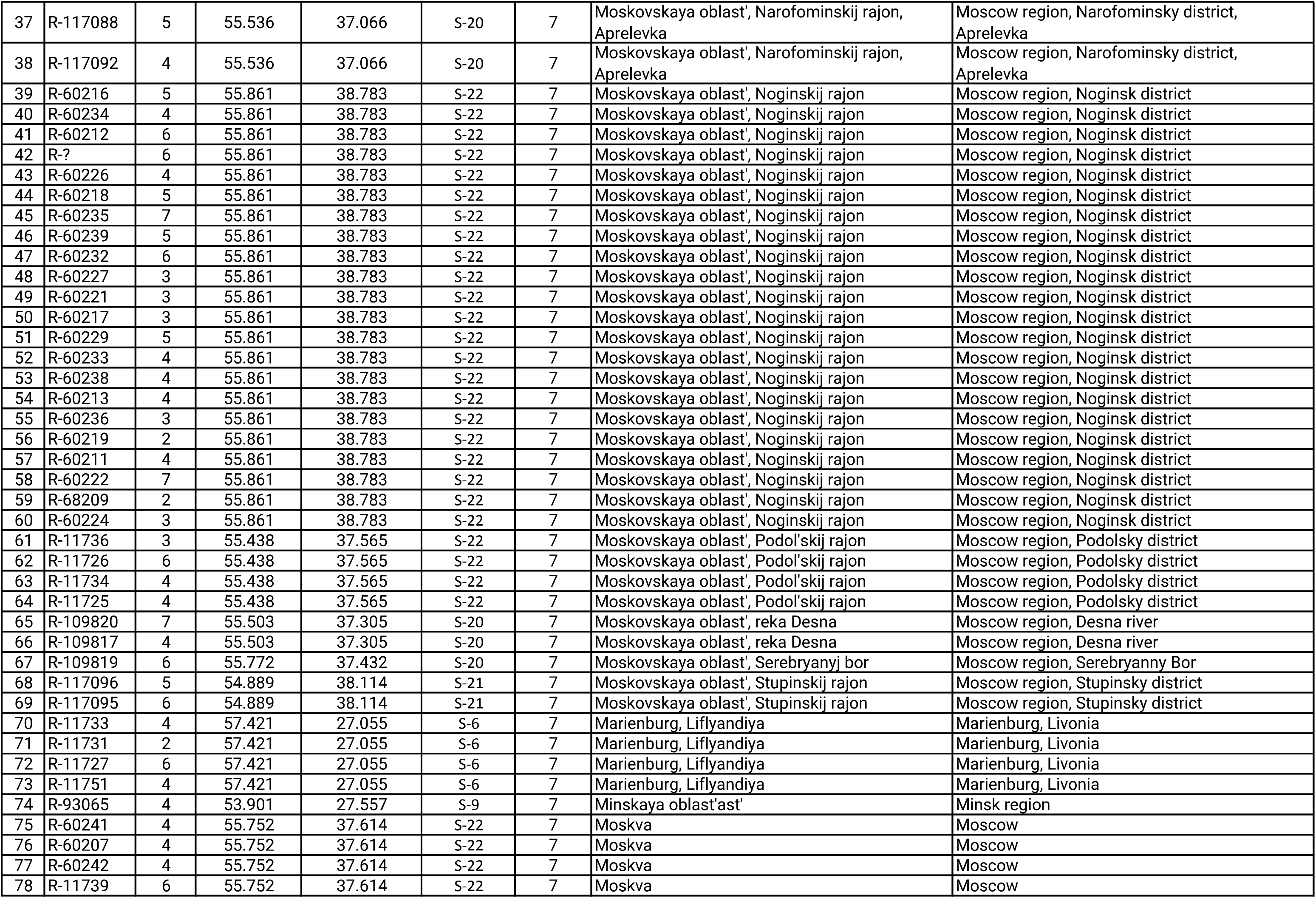
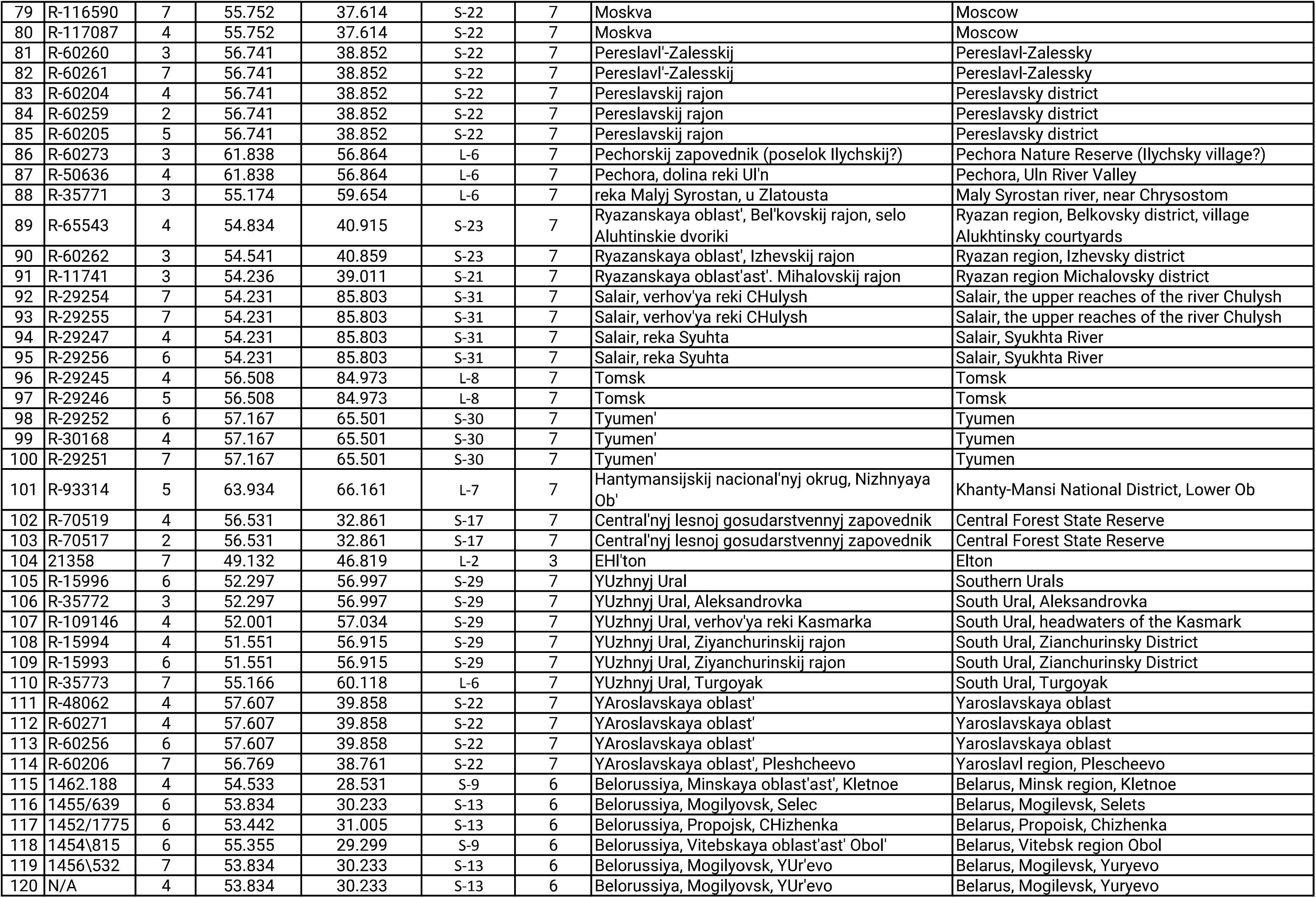
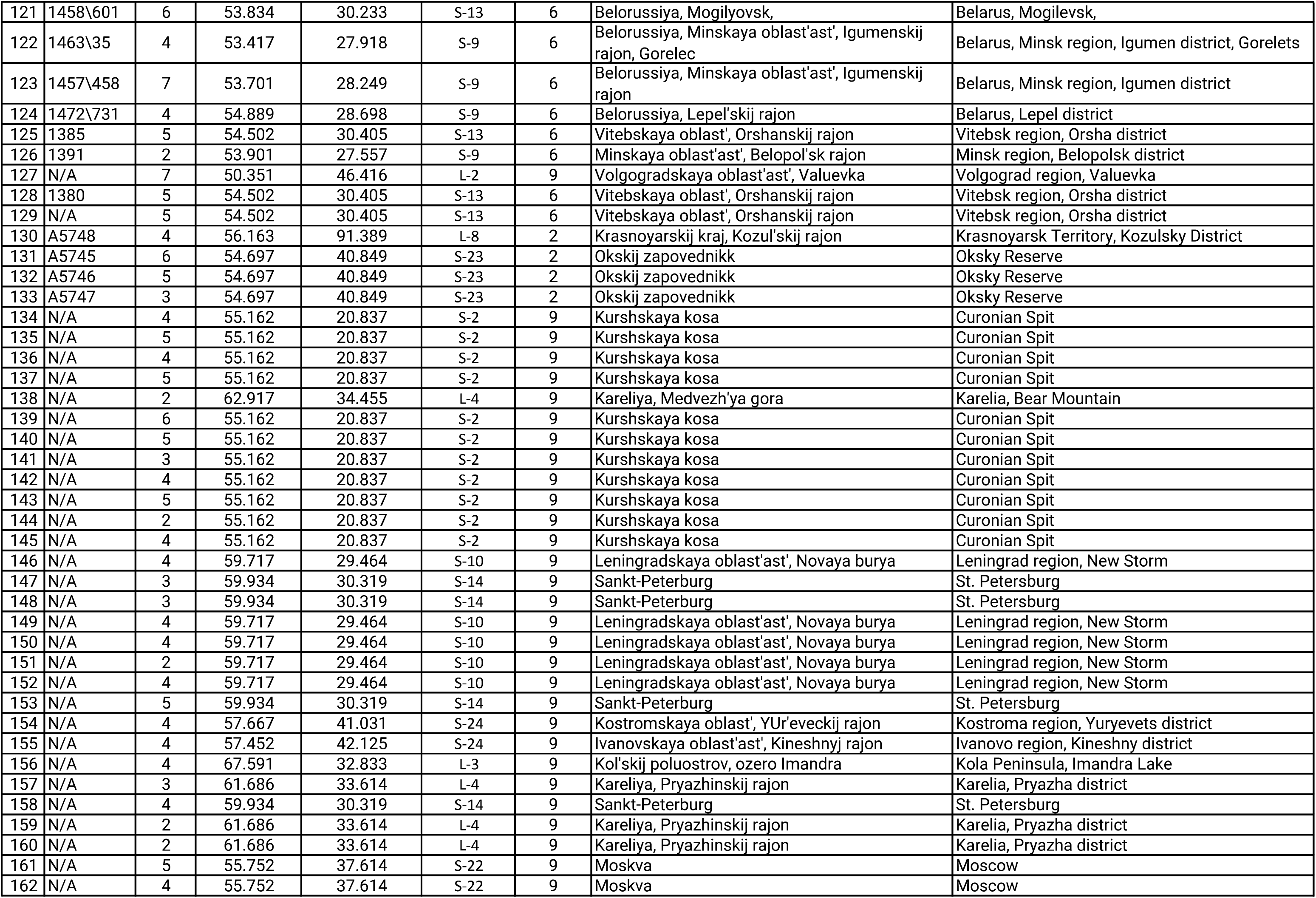
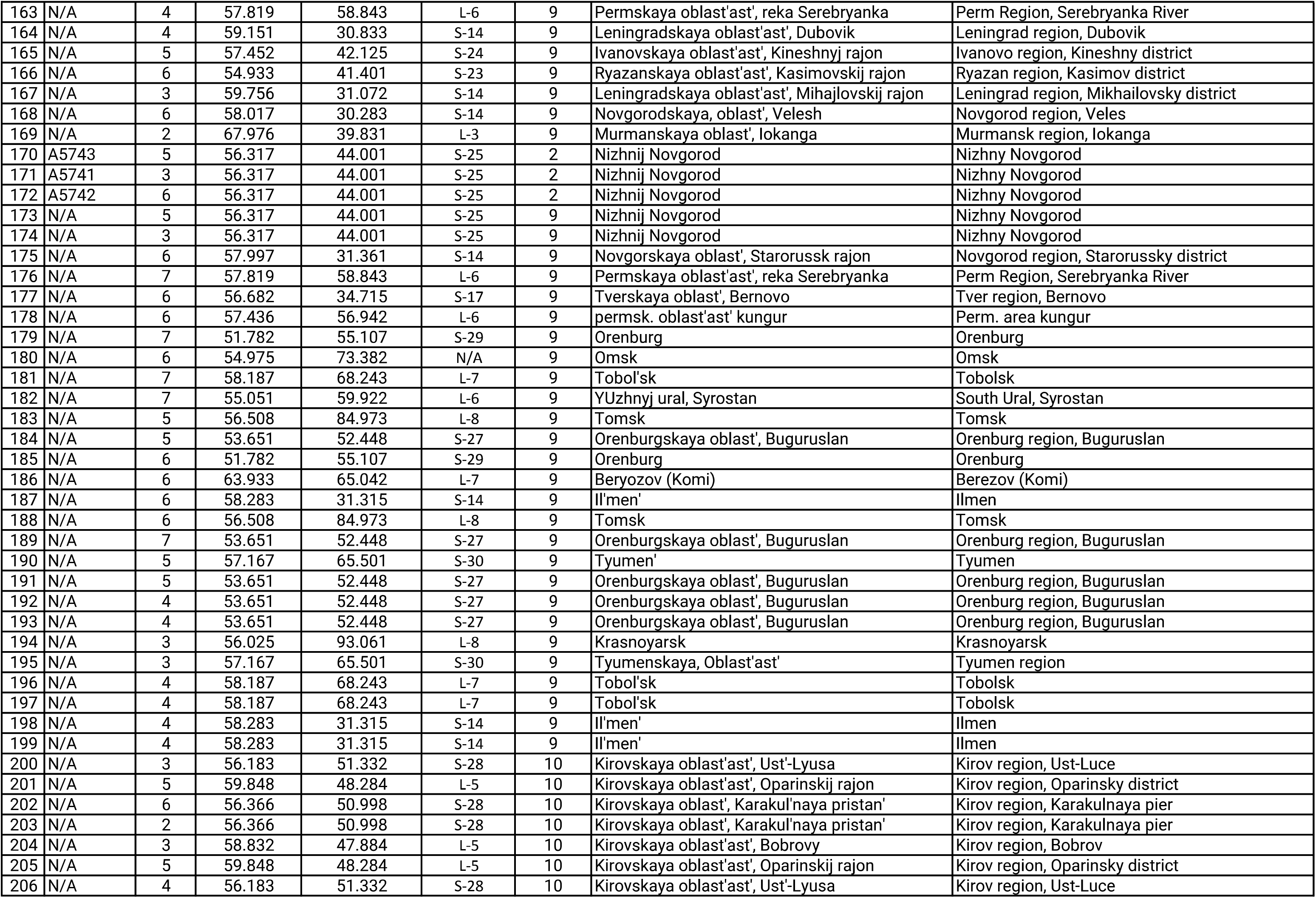
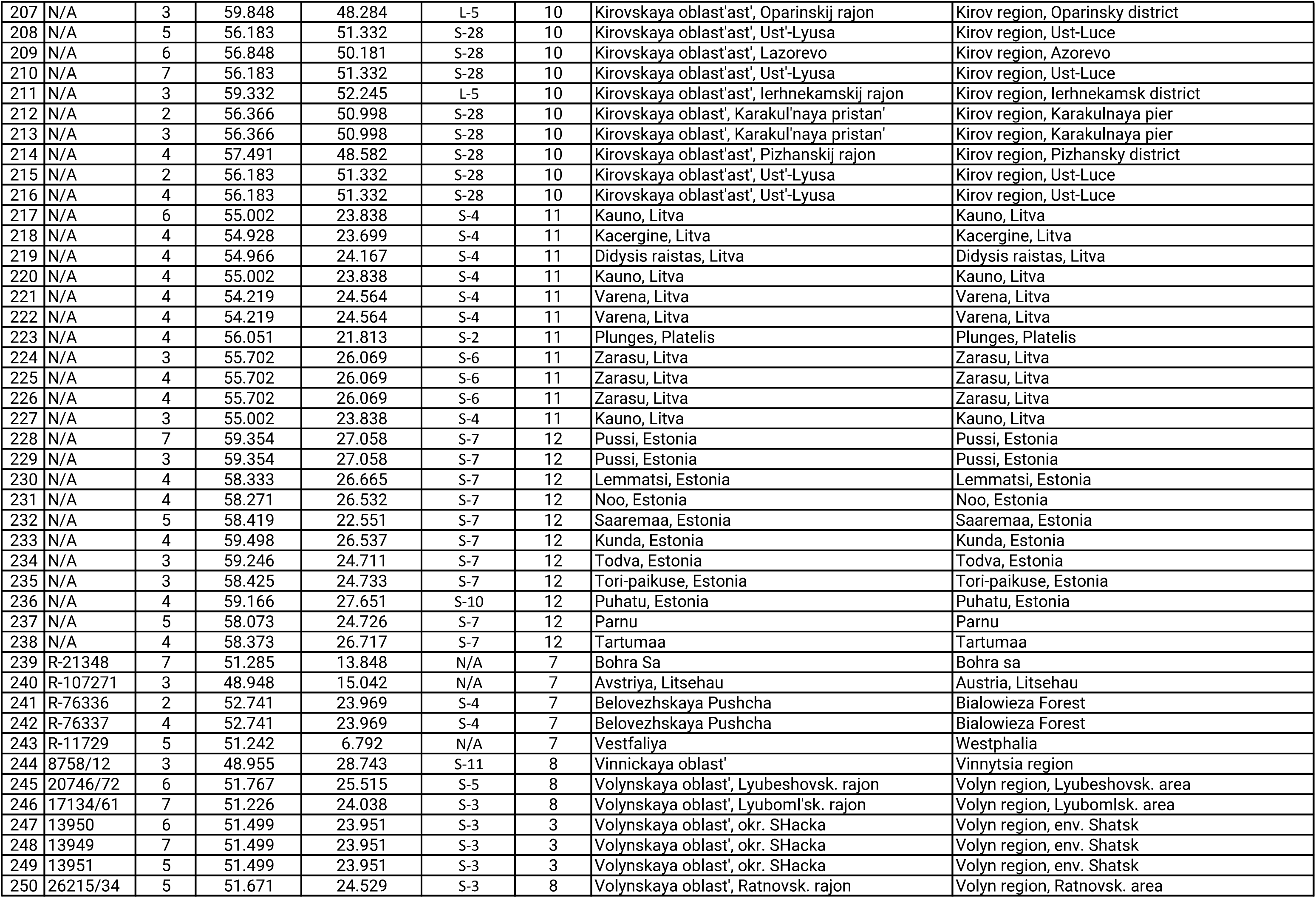
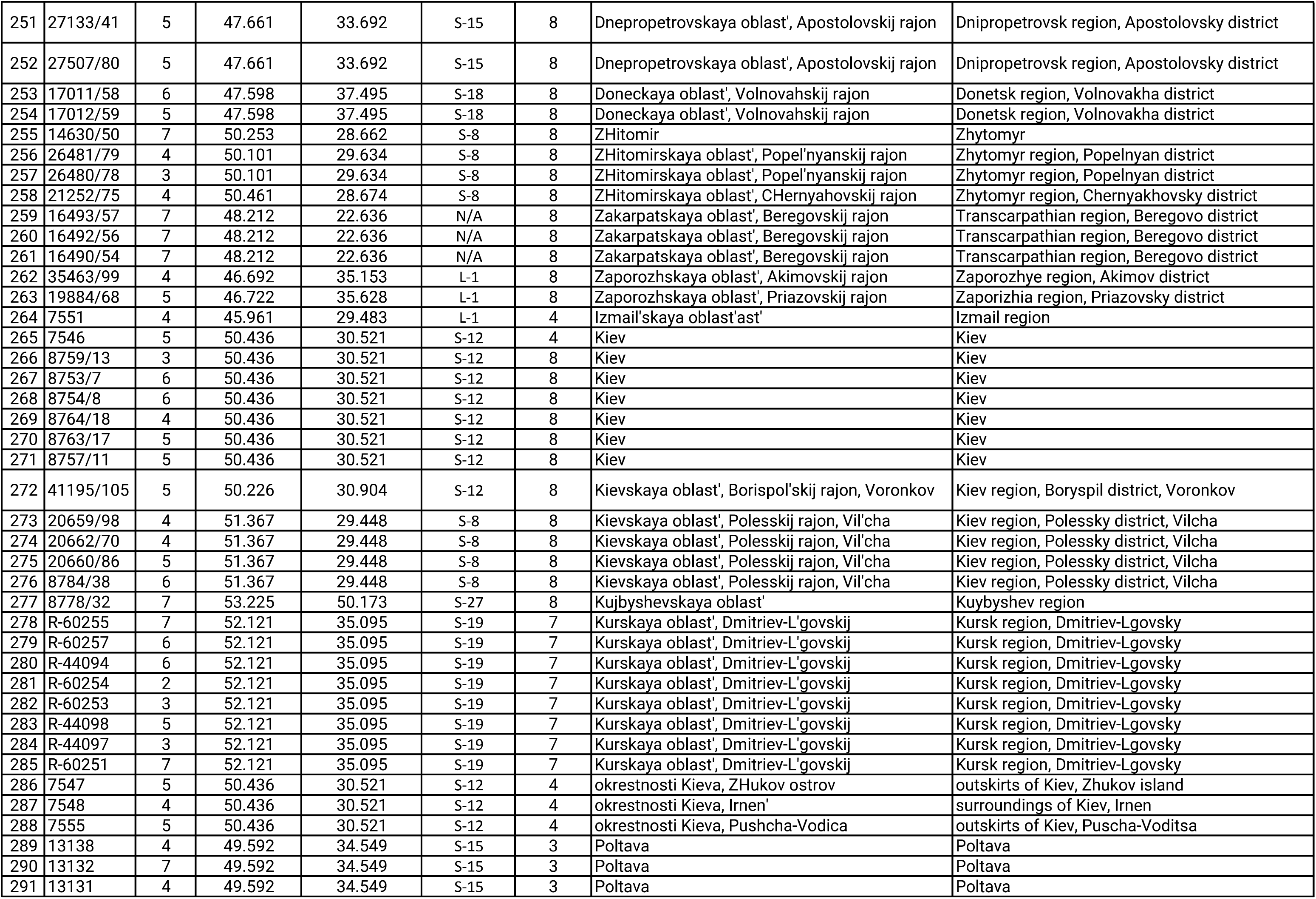
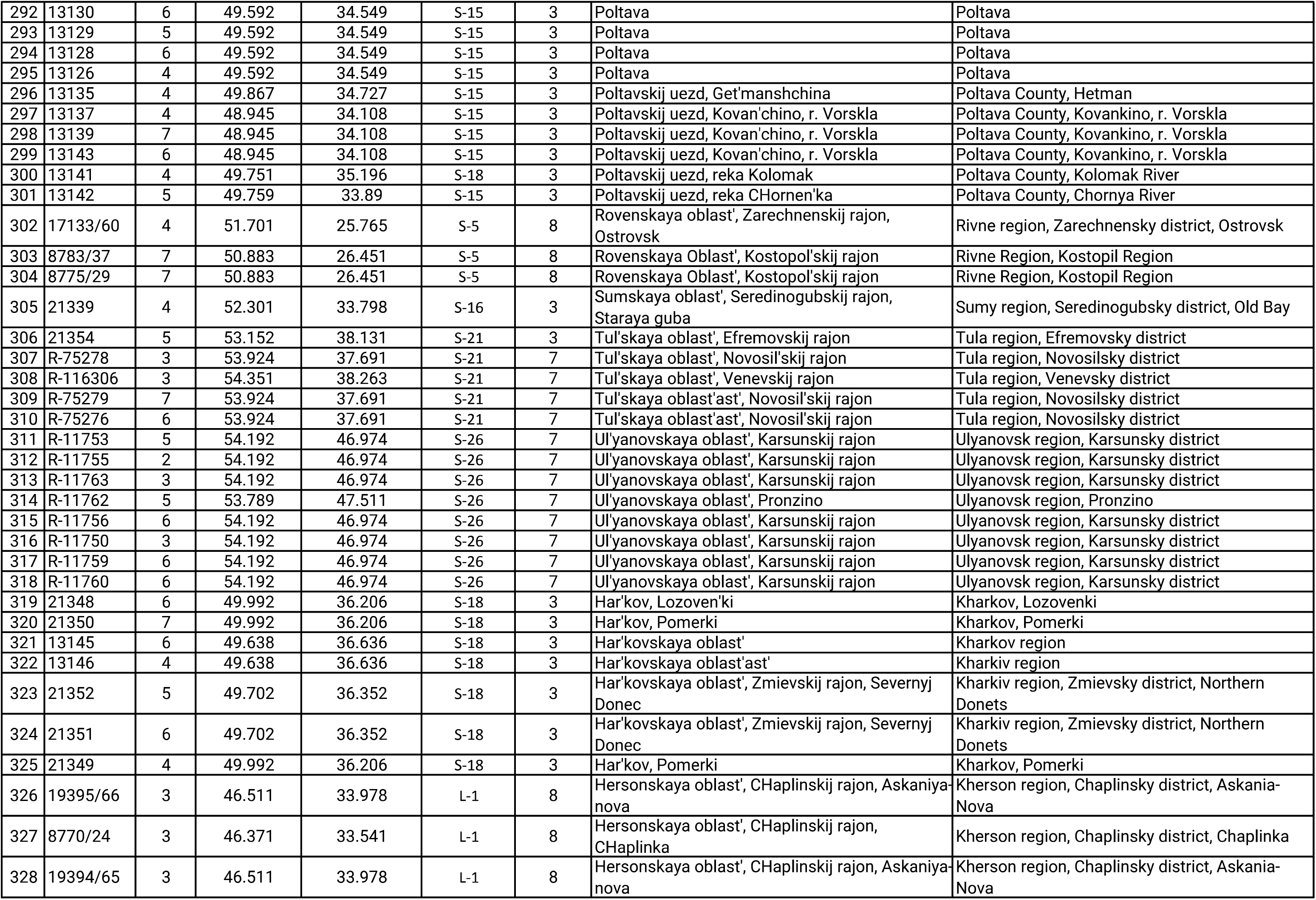
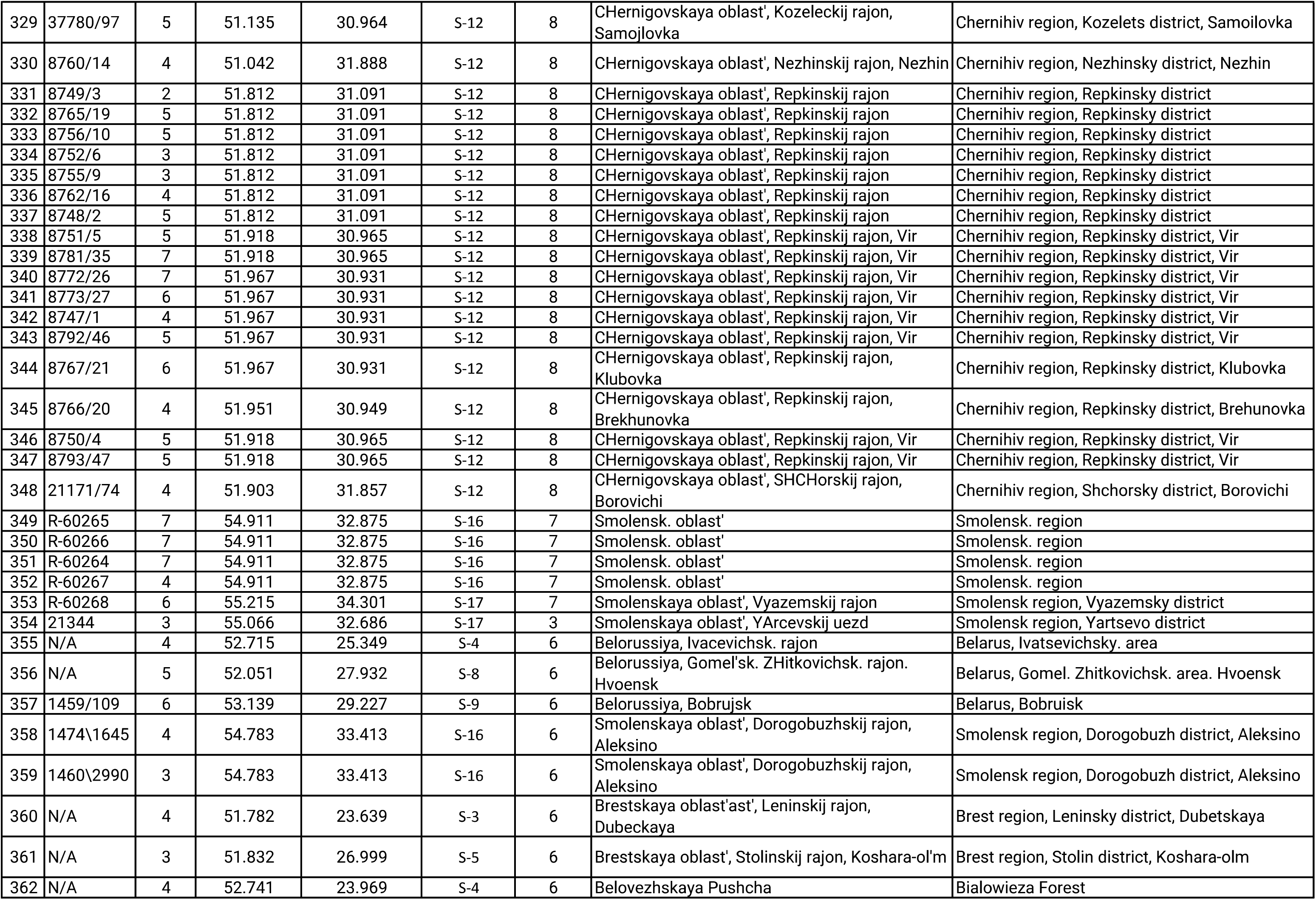
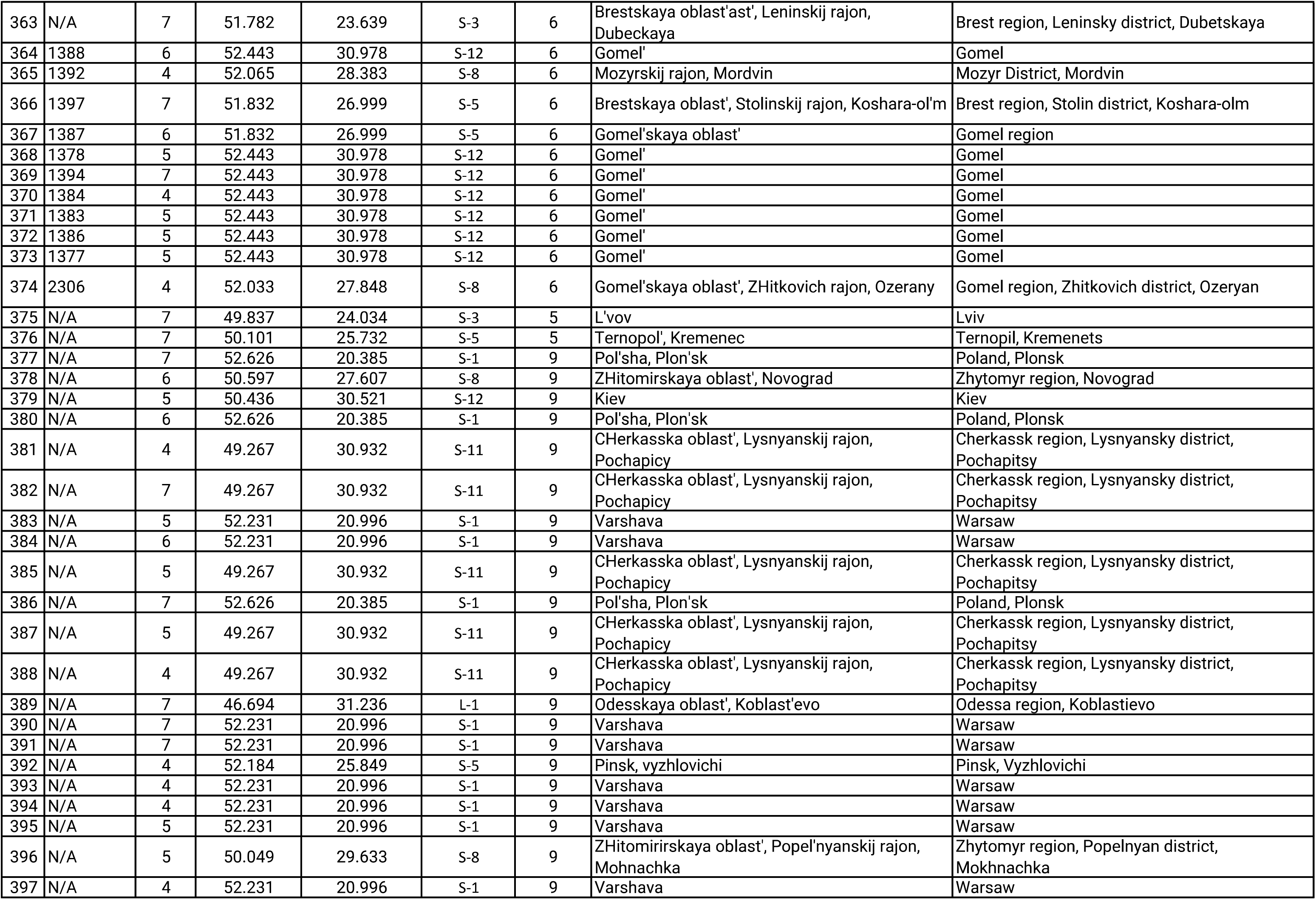
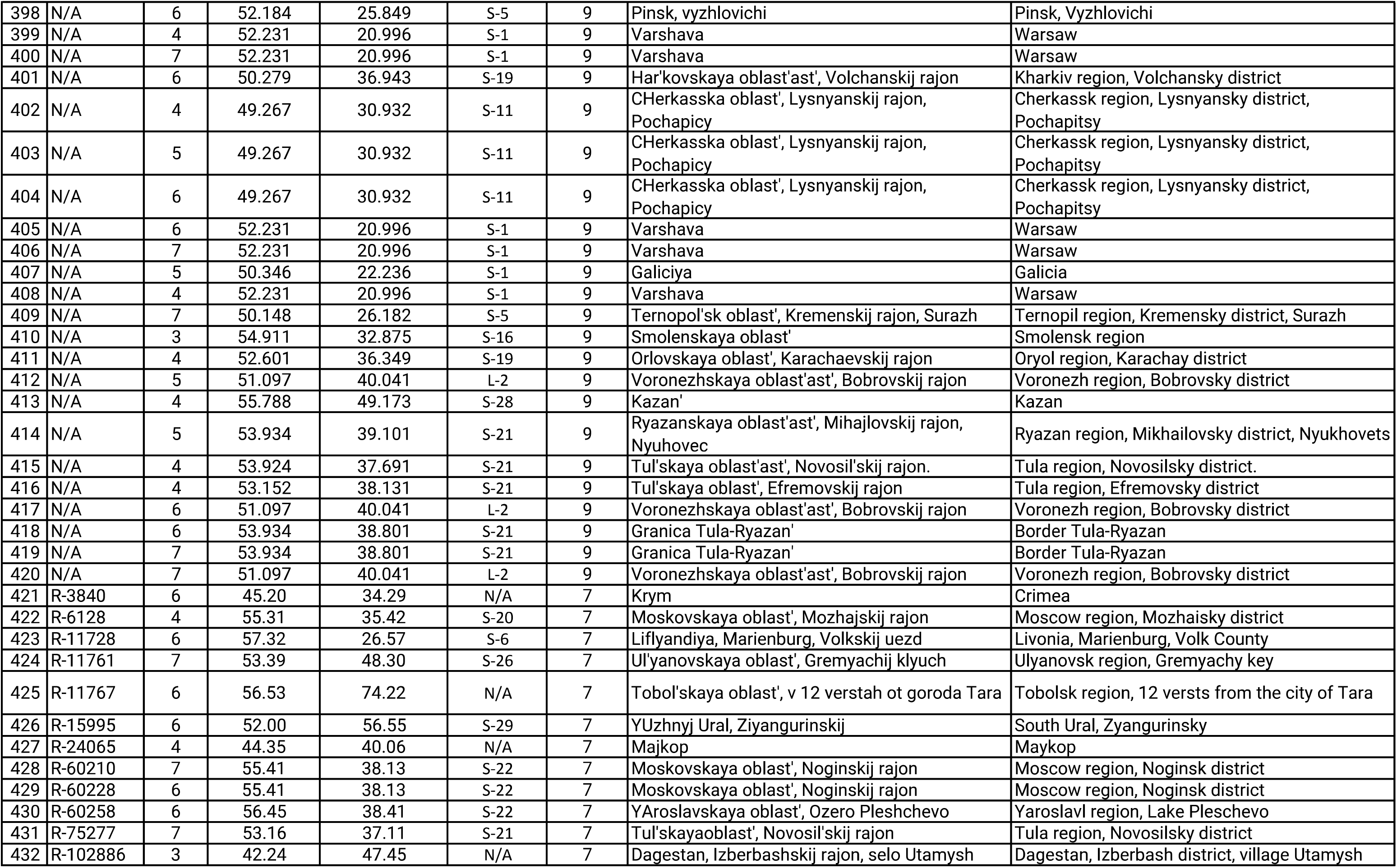
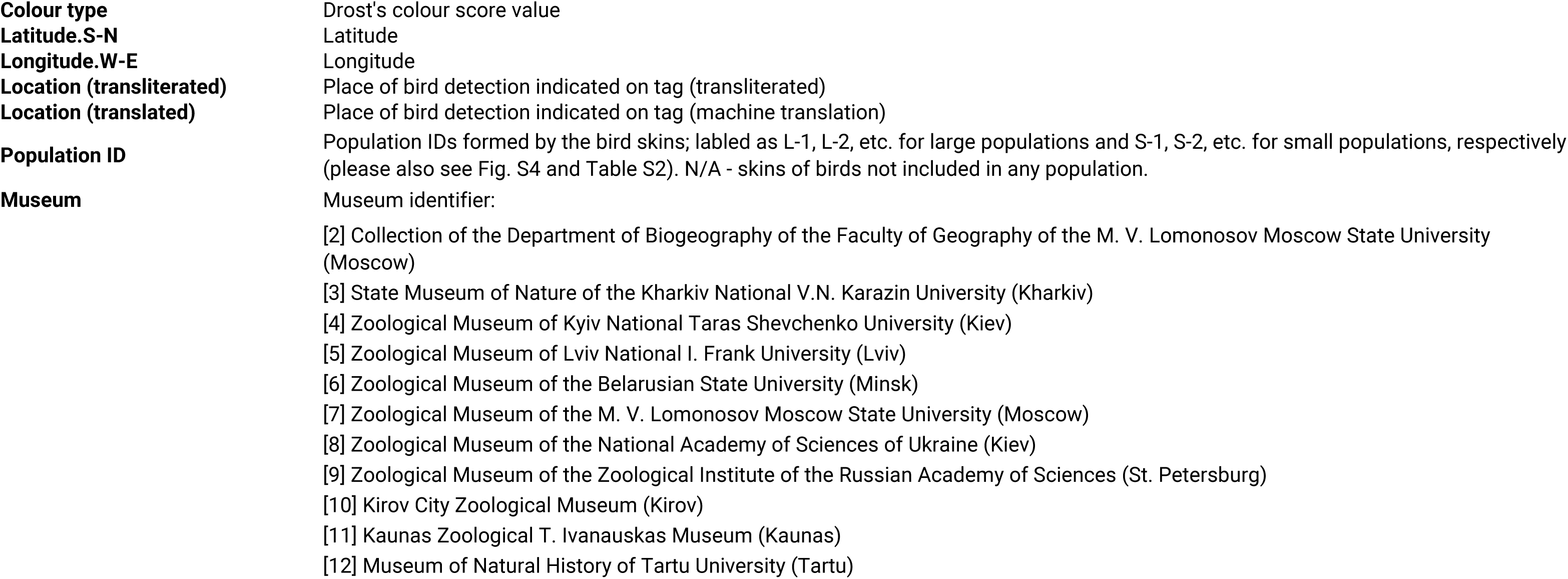
The list of bird skins stored in museum collections and used to calculate the mean breeding plumage color score of males in the Pied Flycatcher populations.

